# Lifestyle Risk Score for aggregating multiple lifestyle factors: Handling missingness of individual lifestyle components in meta-analysis of gene-by-lifestyle interactions

**DOI:** 10.1101/2020.05.26.116723

**Authors:** Hanfei Xu, Karen Schwander, Michael R Brown, Wenyi Wang, RJ Waken, Eric Boerwinkle, L Adrienne Cupples, Lisa de las Fuentes, Diana van Heemst, Oyomoare Osazuwa-Peters, Paul S de Vries, Ko Willems van Dijk, Yun Ju Sung, Xiaoyu Zhang, Alanna C Morrison, DC Rao, Raymond Noordam, Ching-Ti Liu

## Abstract

Recent studies consider lifestyle risk score (LRS), an aggregation of multiple lifestyle exposures, in identifying association of gene-lifestyle interaction with disease traits. However, not all cohorts have data on all lifestyle factors, leading to increased heterogeneity in the environmental exposure in collaborative meta-analyses. We compared and evaluated four approaches (Naïve, Safe, Complete and Moderator Approaches) to handle the missingness in LRS-stratified meta-analyses under various scenarios. Compared to “benchmark” results with all lifestyle factors available for all cohorts, the Complete Approach, which included only cohorts with all lifestyle components, was underpowered, and the Naïve Approach, which utilized all available data and ignored the missingness, was slightly liberal. The Safe Approach, which used all data in LRS-exposed group and only included cohorts with all lifestyle factors available in the LRS-unexposed group, and the Moderator Approach, which handled missingness via moderator meta-regression, were both slightly conservative and yielded almost identical p-values. We also evaluated the performance of the Safe Approach under different scenarios. We observed that the larger the proportion of cohorts without missingness included, the more accurate the results compared to “benchmark” results. In conclusion, we generally recommend the Safe Approach to handle heterogeneity in the LRS based genome-wide interaction meta-analyses.

## 1. Introduction

Thanks to strong collaborations, many large-scale genome-wide association studies (GWAS) have successfully identified many genetic determinants described to explain part of the pathophysiological mechanism underlying a wide range of traits. Despite these efforts and increased sample sizes, the explained variability of many traits is relatively small and only a small proportion of the familial heritability can be explained by the candidate variants found (Evangelou et al., 2018; López-Cortegano & Caballero, 2019; Manolio et al., 2009).

In addition to genetics, environmental factors and gene-environment interactions may contribute to this unexplained trait heritability (Manolio et al., 2009; Rao et al., 2017). Recently, genome-wide gene-environment interaction studies have been conducted to further explore the potential mechanisms underlying an array of diseases or disease traits of interest (de las Fuentes et al., 2020; Graff et al., 2017; Liu et al., 2012; Noordam et al., 2019; Wu et al., 2020). Thus far, these collaborative efforts have largely focused on a single environmental or lifestyle factor, such as smoking (Bentley et al., 2019; Justice et al., 2017; Yun J. Sung et al., 2018; Yun Ju Sung et al., 2019; Wu et al., 2020), physical activity (Graff et al., 2017; Kilpeläinen et al., 2019), alcohol intake (De Vries et al., 2019), educational attainment (de las Fuentes et al., 2020) and others (Jiang et al., 2018; Noordam et al., 2019). By accounting for the environmental risk factor, these efforts identified several novel loci beyond those identified by the traditional main effects-only GWAS. However, multiple environmental factors may simultaneously modify the genetics effects of loci (Osazuwa-Peters et al., 2020). Additionally, single lifestyle variables may not capture the spectrum of relevant environmental variation, resulting in biased effect estimation and false-negative results due to reduced statistical power.

Lifestyle factors, such as smoking, physical inactivity and alcohol consumption, all contribute independently to the risk of developing cardiovascular diseases, and composite lifestyle risk scores (LRS) have been used previously to assess the combined effect of multiple lifestyle factors on cardiovascular disease development (Abdullah Said, Verweij, & Van Der Harst, 2018; Lévesque, Poirier, Després, & Alméras, 2017; Sotos-Prieto, Baylin, Campos, Qi, & Mattei, 2016). However, when applying LRS methodology to large collaborative consortium settings, challenges arise as not all lifestyle components in the LRS are available in all participating cohorts and/or may not be measured using the same instrument. If ignored, significant measurement error and potential heterogeneity may be introduced with reduced statistical power and potential bias. In the present study, we explore different approaches for incorporating missingness of individual lifestyle components with meta-analysis of genome-wide gene-environment interaction on systolic blood pressure in four European-ancestry (EA) cohorts.

## 2. Methods

### 2.1 Participating cohorts and subject inclusion

In this study, we included data from four cohorts, which were the Atherosclerosis Risk in Communities Study (ARIC), the Framingham Heart Study (FHS), the Hypertension Genetic Epidemiology Network (HyperGEN), and the Netherlands Epidemiology of Obesity Study (NEO). For cohorts with data collected from multiple center visits, we chose a single visit that could maximize sample size with non-missing data. We included a total of 24,048 EA individuals who were aged 18-80 and had non-missing genotype, phenotype and relevant covariates information, including age, sex, systolic blood pressure (SBP), anti-hypertensive medications, body mass index (BMI) and the four lifestyle factors (smoking status, alcohol consumption, education level, and physical activity).

### 2.2 Phenotype and covariates

Resting SBP (mmHg) was calculated by taking the average of all available BP readings at the same clinical visit, and further adjusted by adding 15 mmHg for subjects with anti-hypertensive medication use (Tobin, Sheehan, Scurrah, & Burton, 2005). SBP values that were more than six standard deviations away from the mean were winsorized to exactly at six standard deviations from the mean, in order to reduce the potential influence of outliers.

Other covariates included age, sex, field center (if appropriate), and principal components to account for population stratification. Analyses were performed with and without further adjusting for BMI.

### 2.3 Genotyping and QC

Genotyping was performed separately within each cohort using Affymetrix (Santa Clara, CA, USA) or Illumina (San Diego, CA, USA) genotyping arrays (Supplementary Table S1). Each cohort performed imputations with IMPUTE2 (Howie, Donnelly, & Marchini, 2009) or MaCH (Li, Willer, Ding, Scheet, & Abecasis, 2010), using the cosmopolitan reference panel from the 1000 Genomes Project Phase 1 Integrated Release Version 3 Haplotypes (2010-11 data freeze, 2012-03-14 haplotypes) (Altshuler et al., 2012). SNPs were excluded if they were non-autosomal, had minor allele frequency (MAF) <1% or low imputation quality (r^2^<0.1). We conduct further quality control filters centrally during the meta-analysis (Section 2.5.2).

### 2.4 Lifestyle Risk Score

In this study, we considered four lifestyle factors: smoking status (never/former/current smoker), current alcohol intake (drinks per week), educational attainment beyond high school (none/some college/college degree) and physical activity (inactive/active). We classified participants as “college degree” if they completed at least a 4-year college degree, as “some college” if they received any education beyond high school including vocational school but did not complete a college degree and as “none” if they received no education beyond high school (de las Fuentes et al., 2020). Physical activity is expressed in metabolic equivalents (MET; 1 MET = 1 kcal/kg/hour). Inactive individuals were defined as those with <225 MET-minutes per week of moderate-to-vigorous leisure-time or commuting physical activity, or in the lower quartile (25%) of the physical activity distribution within cohort. The detailed definitions of active and inactive physical activity followed a previous study on gene-physical activity interaction (Kilpeläinen et al., 2019).

Construction of the lifestyle risk score can be separated into two steps. First, each lifestyle factor, treated as an individual lifestyle component, was categorized into no risk (with value of 0), low risk (with value of 1) and high risk (with value of 2) based on its effect on BP or cardiovascular health, except physical activity which only had no risk and low risk (Osazuwa-Peters et al., 2020). The higher risk value the category was assigned, the more relevant to unfavorable cardiovascular health outcomes. Note that we categorized modest alcohol intake as no risk and abstinence as low risk because there was evidence that moderate alcohol consumption had consistently been associated with a decreased risk of type 2 diabetes (Joosten et al., 2010) and coronary artery disease (Klatsky, 1999) compared with abstention or excessive consumption (Feitosa et al., 2018). Table 1 provides the details of lifestyle risk score component definition.

**Table 1.** Definition of Lifestyle Risk Score Component, with no risk as the value of 0, low risk as the value of 1 and high risk as the value of 2.

| Component variables | No risk (0) | Low risk (1) | High risk (2) |
| --- | --- | --- | --- |
| Smoking | Never | Former | Current |
| Current Alcohol Intake (drinks/week) | Modest (1 – 7) | Abstinence* (0) | Heavy (> 7) |
| Education (after High School) | College Degree | Some College | None |
| Physical Activity | Active | Inactive | -- |

*\* Includes Former Drinkers*

Second, the “Complete” Quantitative Lifestyle Risk Score (QLRS-C) was calculated by summing up all four components, ranging from 0-7. We also calculated the “Partially Missing” Quantitative Lifestyle Risk Score (QLRS-M) using 2-3 components pre-selected for each cohort by design, as described in Table 2. For example, for ARIC, we included three lifestyle components (smoking, education and physical activity) when constructing QLRS-M. QLRS-M ranges from 0 to 4 or 5, depending on the inclusion of lifestyle components for each cohort.

**Table 2.** Components included in the calculation of “Partially Missing” Lifestyle Risk Score (QLRS-M) for each cohort (by design)

| Cohort | Smoking | Alcohol | Education | Physical Activity |
| --- | --- | --- | --- | --- |
| ARIC | Include | Not Include | Include | Include |
| FHS | Not Include | Include | Include | Include |
| HyperGEN | Not Include | Include | Include | Not Include |
| NEO | Include | Not Include | Include | Not Include |

After constructing the Quantitative Lifestyle Risk Scores, we further created Dichotomous Lifestyle Risk Scores for the “Complete” (DLRS-C) and the “Partial” (DLRS-M) summary scores. We gave a value of 0 (unexposed group) if the corresponding Quantitative Lifestyle Risk Score <2 and a value of 1 (exposed group) if Quantitative Lifestyle Risk Score ≥2 (i.e. at least one risk component classified as high risk or at least two components classified as low risk). These dichotomized LRS measures are used to define exposed and unexposed strata in our analyses.

It is worth noting that cohorts with partially missing lifestyle components have equal or lower LRS than its actual score had we observed all lifestyle components. This leads to potential misclassification when dichotomizing the LRS into exposed and unexposed groups. However, no participant would be misclassified as exposed and they can only be misclassified as unexposed, leading to heterogeneity in the unexposed group only.

### 2.5 Statistical Analysis

#### 2.5.1 Overview

We conduct a two-stage analysis procedure. In Stage 1, each cohort performed LRS-stratified genome-wide association analysis on SBP using the main effect model (*E*(*Y*) = *β_0_* + *β_G_ SNP* + *β_C_ Covariates*, where *Y* is the SBP level, *SNP* is the imputed additive dosage value of the genetic variant), in DLRS-C exposed and DLRS-C unexposed strata. The association analyses were also repeated in DLRS-M exposed and DLRS-M unexposed strata. In Stage 2, we performed meta-analysis within each stratum, and then evaluated the joint effects of main and interaction effects by calculating the p-values for the 2 degree of freedom joint test. Under Stage 2, we considered four different meta-analysis approaches of handling missingness of lifestyle components (Naïve, Safe, Complete and Moderator Approaches). We evaluated the performance of the four approaches under four scenarios where some cohorts were designed to provide association results using “Complete” LRS but the others were designed to only provide “Partially Missing” results.

#### 2.5.2 Stage 1: Cohort-specific stratified analysis and QC of association results

For Stage 1, each cohort performed eight genome-wide association analyses on SBP using the main effect model: two strata (exposed/unexposed) × two LRS (DLRS-C/DLRS-M) × two BMI adjustment (with/without). Association analyses were implemented either using ProbABEL (Aulchenko, Struchalin, & van Duijn, 2010) for studies with unrelated samples, or using MMAP (https://mmap.github.io/) for studies with family relatedness. Each cohort provided the robust estimates of the stratum-specific genetic main effect and corresponding robust standard error (SE) for all eight analyses. Cohort-specific details are presented in Supplementary Table S1.

We performed extensive quality control (QC) using the R package EasyQC (Winkler et al., 2014) on each of the eight cohort-specific association results, which contained approximately 8-9 million variants. First, we removed variants with invalid alleles and indels, harmonized alleles and variant names across cohorts, and compared allele frequencies with the ancestry-specific 1000 Genomes reference panel. Next, we compared summary statistics (e.g., mean, standard deviation, minimum, maximum) of estimated effect sizes, standard errors, and p-values across cohorts to identify potential outliers, and reviewed SE-N (i.e., inverse of the median standard error versus the square root of the sample size) plots to look for possible problems with phenotype or covariates (Winkler et al., 2014). Finally, SNPs were excluded if imputation quality score was <0.5, or if the product of the imputation quality and minor allele count was <20 (de las Fuentes et al., 2020; Kilpeläinen et al., 2019). No genomic control was applied after filtering as there were little to no problems with inflation (genomic control inflation λ ranged from 1.020 to 1.071).

Since QC and filtering were performed separately within each stratum, the set of variants remaining in each stratum differed slightly. Thus, we further harmonized the set of variants between the exposed and unexposed strata within each LRS construction – BMI adjustment combination, to ensure that the set of variants was identical between strata. After QC, the number of variants in each association result was between 5.3M-8.2M.

#### 2.5.3 Stage 2: Meta-analysis

After obtaining cohort-specific GWAS results using “Complete” and “Partially Missing” LRS, we first performed meta-analyses within each stratum (exposed/unexposed) using the results obtained from analyses using “Complete” LRS, and considered this set of meta-analyzed results as the “benchmark” results as there is no missing lifestyle component in each cohort’s LRS construction.

Then, to mimic the real life situation where some of the cohorts would provide GWAS association results obtained from analyses using “Complete” LRS but the others could only provide “Partially Missing” results, we further performed the meta-analyses using a mixture of results obtained from cohort-specific analyses conducted with “Complete” LRS and “Partially Missing” LRS. We considered four scenarios using different cohort mixture patterns by changing each cohort’s contribution of lifestyle components, in order to better utilize the data. The setting of each scenario is presented in Table 3. For example, Scenario 1 is to use “complete” results from ARIC, and “partially missing” results from HyperGEN, FHS and NEO.

**Table 3.** Setting of Scenarios 1-4 using a mixture of results obtained from cohort-specific genome-wide association analyses conducted with “Complete” LRS and “Partially Missing” LRS, and the inclusion of association results in the meta-analysis using Naïve, Safe, Complete and Moderator Approaches.

| Scenario | Strata | Cohort |  |  |  |
| --- | --- | --- | --- | --- | --- |
|  |  | ARIC | FHS | HyperGEN | NEO |
| Scenario 1 | Unexposed | Complete<sup>N,S,C,M</sup> | Partially Missing<sup>N,M</sup> | Partially Missing<sup>N,M</sup> | Partially Missing<sup>N,M</sup> |
|  | Exposed | Complete<sup>N,S,C,M</sup> | Partially Missing<sup>N,S,M</sup> | Partially Missing<sup>N,S,M</sup> | Partially Missing<sup>N,S,M</sup> |
| Scenario 2 | Unexposed | Partially Missing<sup>N,M</sup> | Complete<sup>N,S,C,M</sup> | Partially Missing<sup>N,M</sup> | Partially Missing<sup>N,M</sup> |
|  | Exposed | Partially Missing<sup>N,S,M</sup> | Complete<sup>N,S,C,M</sup> | Partially Missing<sup>N,S,M</sup> | Partially Missing<sup>N,S,M</sup> |
| Scenario 3 | Unexposed | Complete<sup>N,S,C,M</sup> | Partially Missing<sup>N,M</sup> | Partially Missing<sup>N,M</sup> | Complete<sup>N,S,C,M</sup> |
|  | Exposed | Complete<sup>N,S,C,M</sup> | Partially Missing<sup>N,S,M</sup> | Partially Missing<sup>N,S,M</sup> | Complete<sup>N,S,C,M</sup> |
| Scenario 4 | Unexposed | Complete<sup>N,S,C,M</sup> | Partially Missing<sup>N,M</sup> | Complete<sup>N,S,C,M</sup> | Complete<sup>N,S,C,M</sup> |
|  | Exposed | Complete<sup>N,S,C,M</sup> | Partially Missing<sup>N,S,M</sup> | Complete<sup>N,S,C,M</sup> | Complete<sup>N,S,C,M</sup> |

<sup>N</sup>: included in the meta-analysis using Naïve Approach; <sup>S</sup>: included in the meta-analysis using Safe Approach; <sup>C</sup>: included in the meta-analysis using Complete Approach; <sup>M</sup>: included in the meta-analysis using Moderator Approach.

As mentioned in the LRS section, the missingness in lifestyle components will cause misclassification when dichotomizing LRS into exposed and unexposed groups, hence leading to heterogeneity in the unexposed group only. To account for this heterogeneity, we considered four different meta-analysis approaches of utilizing “Complete” and “Partially Missing” results under various scenarios discussed above.

1. Naïve Approach. This approach simply takes all association results contributed by each participating cohort without worrying whether their LRS includes all lifestyle components, for both exposed and unexposed groups.
2. Safe Approach. Since heterogeneity only occurs in the unexposed group, it is “safe” to only take association results from cohorts with complete LRS for the unexposed group analysis, while including results from all cohorts no matter whether the missing data exist in LRS for the exposed group analysis.
3. Complete Approach. This approach only uses association results from cohorts with complete LRS data in meta-analysis, for both exposed and unexposed groups.
4. Moderator Approach. This approach uses all the contributed data from cohorts without regard to their missingness in lifestyle components. It utilizes the framework of meta-regression, while including moderator terms indicating the missing LRS components across cohorts in the design matrix of the meta-regression to account for missingness during meta-analysis. Technical details of this approach are available in the Supplementary Method.

Table 3 also shows the inclusion of association results in the meta-analysis using each of the approaches described above under Scenarios 1-4. Here we take Scenario 1 as an example: For the Naïve Approach, we analyze exposed and unexposed groups separately using “complete” results from ARIC, and “partially missing” results from HyperGEN, FHS and NEO without differentiating “complete” or “partially missing”. For the Safe Approach, we include ARIC results alone and ignore other cohorts’ contributions with “partially missing” results for the unexposed group; for the exposed group, we analyze all four cohorts using “complete” results from ARIC, and “partially missing” results from HyperGEN, FHS and NEO. For the Complete Approach, we analyze exposed and unexposed groups separately, but only use “complete” results from ARIC with no other cohorts included. For the Moderator Approach, we take “complete” results from ARIC, and “partially missing” results from HyperGEN, FHS and NEO for both exposed and unexposed groups as input of the meta-regression.

For the “benchmark” meta-analysis and the first three approaches (Naïve, Safe and Complete), we used METAL software (Willer, Li, & Abecasis, 2010) to perform meta-analyses within each stratum and used EasyStrata (Winkler et al., 2015) to calculate the 2 degree of freedom joint p-values. For the Moderator Approach, we used the Moderator Web App and R code developed by Dr. RJ Waken (https://rjwaken.shinyapps.io/missing_lrs_meta/).

In the following sections, we focus on the comparison of results obtained from the analyses without adjusting for BMI, since we observed the same pattern for BMI-adjusted analyses and the primary objective of our study is to evaluate the meta-analysis approaches of handling missingness instead of identifying novel loci under confounder adjustment.

## 3. Results

### 3.1 Sample Characteristics

Sample characteristics are presented in Supplementary Tables S2, S3, and S4. ARIC had the largest sample size (N=9,426) and HyperGen cohort had the fewest number of participants (N=1,249). All cohorts had similar distributions of sex, age and BMI, except that FHS and HyperGEN had a wider age range than ARIC and NEO (Supplementary Table S2). In Supplementary Tables S3 and S4, the exposed group had slightly higher SBP level than the unexposed group for all four cohorts in terms of DLRS-C. However, the difference in SBP levels between exposed and unexposed groups was smaller when we defining exposure groups using DLRS-M. The proportion of subjects in the exposed group was smaller when using DLRS-M compared to DLRS-C, indicating potential misclassification.

### 3.2 Results Comparison between Approaches

Figure 1 presents the results of comparison of the four meta-analysis approaches to the “benchmark” results derived from analyses of cohorts where all lifestyle components were present. Among variants that reach genome-wide significance level (p-value<5×10^−8^), we observed that the Complete Approach yielded much larger p-values than the “benchmark” results, thus could be considered underpowered. The Naïve Approach was able to detect the same set of genome-wide significant variants as the “benchmark” results, but with slightly smaller p-values. The Safe and Moderator Approaches led to slightly larger p-values than “benchmark” results. The Q-Q plot (Figure 2) also shows that the Complete Approach obtained the most deflated p-values among the four approaches (λ_Complete vs benchmark_ = 0.972). The Safe Approach and Moderator Approach yielded similar slightly conservative results (λ_Safe vs benchmark_ = λ_Moderator vs benchmark_ =0.985), while the results of the Naïve Approach were slightly liberal (λ_Naive vs benchmark_ = 1.004).

**Figure 1.**
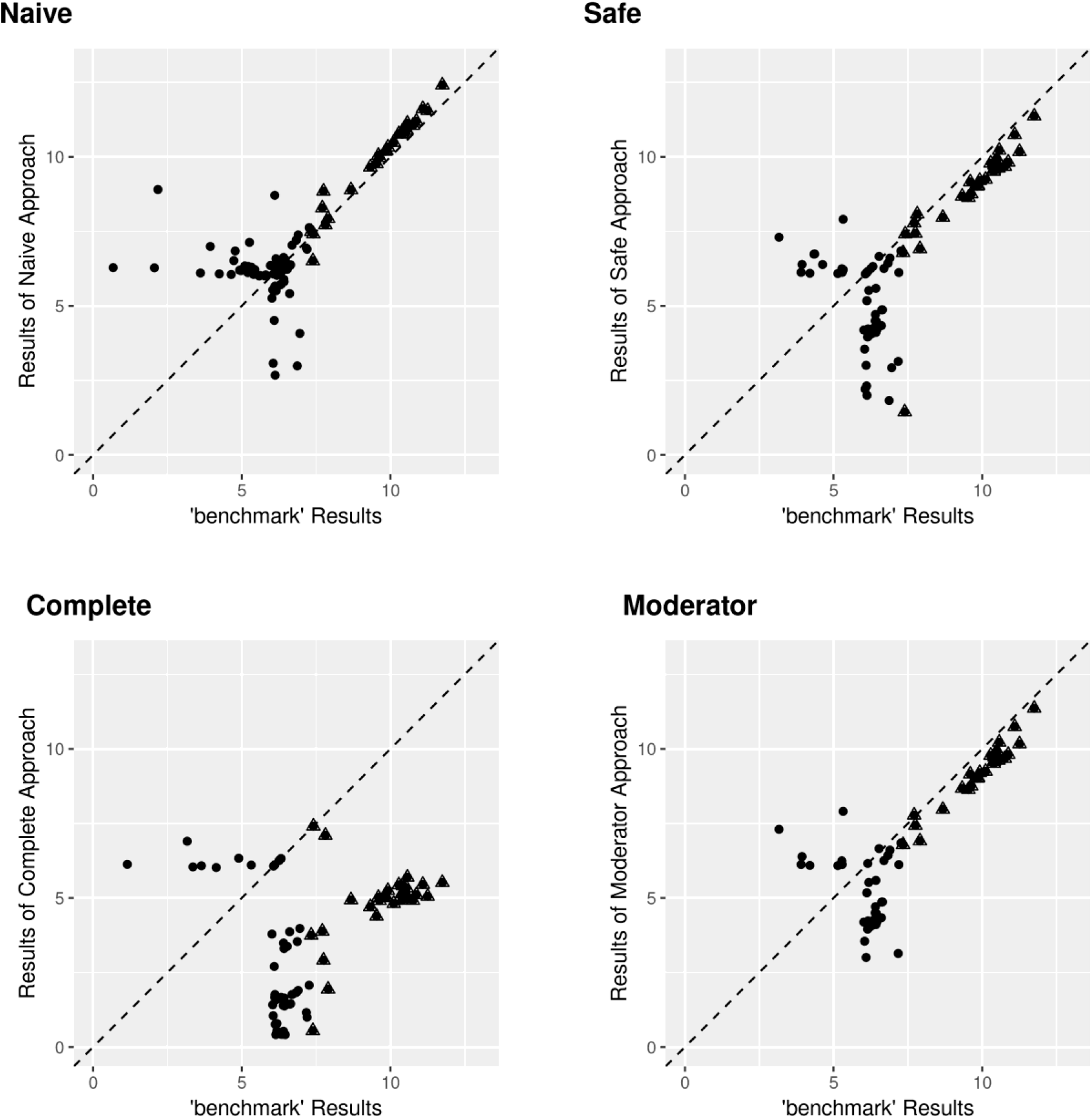
Scatterplots of comparison of four approaches to “benchmark” results in terms of – log_10_ (p-value). Each plot shows SNPs with p-value<10^−6^ for any of the two approaches being compared in the plot. SNPs reaching genome-wide significant (p-value<5×10^−8^) in “benchmark” results are marked as triangle.

**Figure 2:**
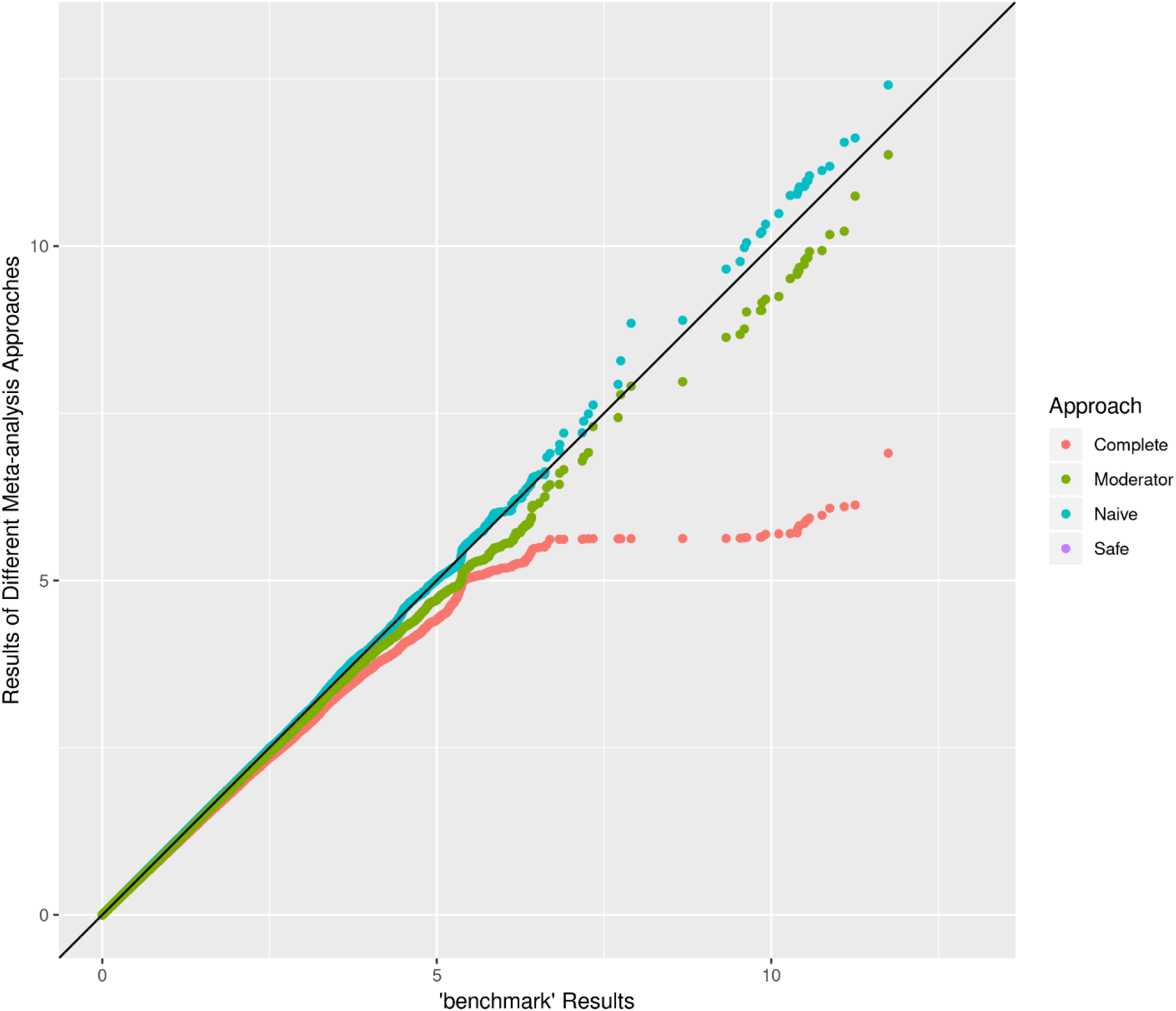
Q-Q plot of different approaches compared to “benchmark” results for Scenario 1 (Scenario 1: use “complete” results from ARIC, and “partially missing” results from HyperGEN, FHS and NEO). λ_Naive_ = 1.004, λ_Safe_ = λ_Moderator_ =0.985, λ_Complete_ = 0.972.

Figure 3 shows the pair-wise comparison among four meta-analysis approaches. Similar to what we observed in Figure 1, the Complete Approach was underpowered compared to other three approaches. The Safe and Moderator Approaches yielded similar but slightly larger p-values than Naïve Approach, and the degree of similarity increased with significance. Notably, the results of Safe Approach and Moderator Approach were almost identical, but the number of variants included in the analyses for the Moderator Approach (Number of variants = 5,258,666) was much smaller than the Safe Approach (Number of variants =8,181,669).

**Figure 3.**
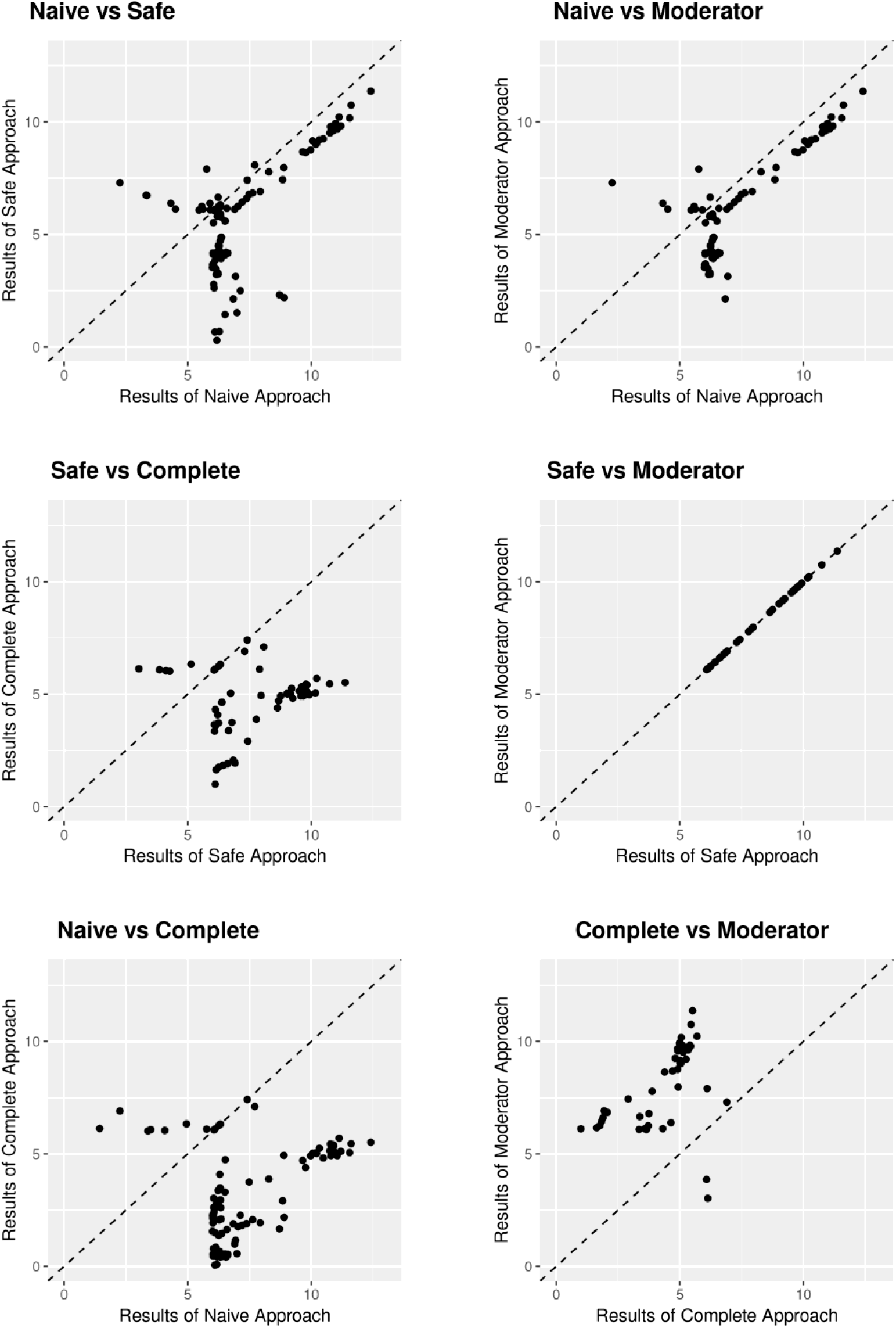
Scatterplots of comparison between four approaches in terms of −log_10_ (p-value). Each plot shows SNPs with p-value<10^−6^ for any of the two approaches being compared in the plot.

Scenarios 2-4 presented similar patterns to Scenario 1 in terms of comparison with “benchmark” results and within-scenario comparison between different approaches; so we focus on illustrating Scenario 1 in the results section. The detailed comparison results of Scenarios 2-4 are available in the Supplementary Figures S1-S9.

### 3.3 Result Comparison between Scenarios for Safe Approach

Here we further evaluated the performance of the same meta-analysis approach under different scenarios. Since we generally were concerned with false-positive results, we focused our attention only to the non-inflated Safe Approach. Figure 4 presents the scatterplot of association results between “benchmark” and the Safe Approach for each of the four scenarios, for variants with p-value<1×10^−6^ in at least one of comparing results. We observed that for SNPs reaching genome-wide significance (p-value<5×10^−8^) in “benchmark” results, the points of Scenarios 3 and 4 almost lay along the diagonal line, while points of Scenarios 1 and 2 were a bit away from the diagonal. This indicated that the Safe Approach under Scenarios 3 and 4 more accurately identified positive signals than under Scenarios 1 and 2.

**Figure 4.**
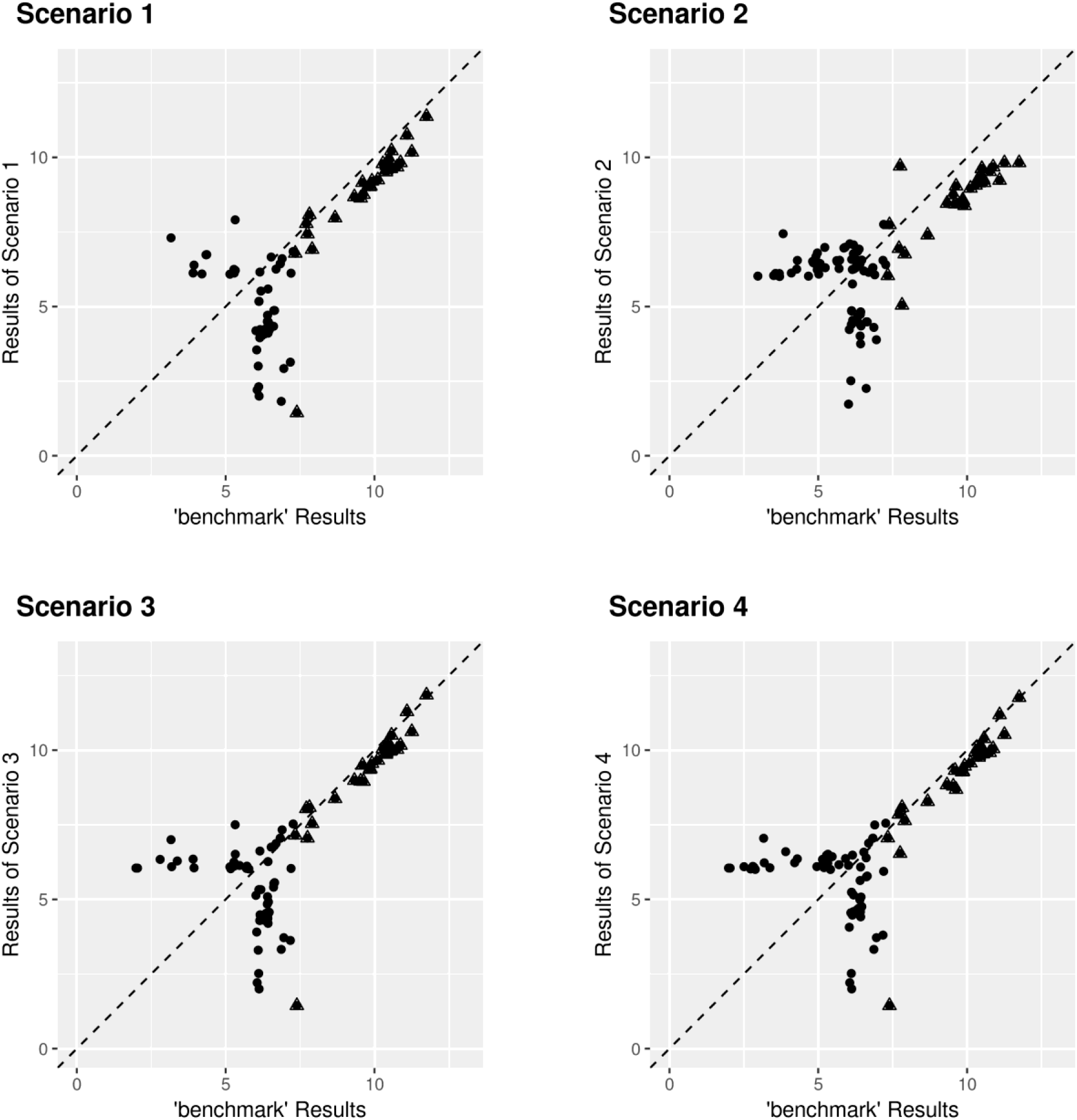
Scatterplots of comparison of four scenarios to “benchmark” results in terms of −log_10_ (p-value) for Safe Approach. Each plot shows SNPs with p-value<10^−6^ for any of the two approaches being compared in the plot. SNPs reaching genome-wide significant (p-value<5×10^−8^) in “benchmark” results are marked as triangle.

The Q-Q plot (Supplementary Figure S10) shows that when p-values were large (>10^−5^), Scenario 4 with less missingness provided more similar p-value distributions with “benchmark” results (λ_scenario 4 vs benchmark_=0.991) compared to Scenario 1 (λ_scenario 1 vs benchmark_=0.983) and Scenario 3 (λ_scenario 3 vs benchmark_=0.984). Although Scenario 2 seemed to perform very well on large p-values (λ_scenario 2 vs benchmark_=0.994), it provided substantially deflated results toward the tail when reaching genome-wide significance. In the meantime, Scenarios 3 and 4 had similar p-value distributions and both of their p-values were very close to the “benchmark” distribution when p-values were small. The p-values of Scenario 1 were closer to the diagonal line than those of Scenario 2 when p-values were small, and this may due to the sample size of the cohort with “complete” results in Scenario 1 (ARIC, N=9,426), which was greater than that of Scenario 2 (FHS, N=7,638).

In general, we consider Scenario 4 performed better than Scenario 3, in turn better than Scenarios 1 and 2. This meets our expectation as Scenario 4 had the smallest proportion of cohorts using “partially missing” results; thus it was expected to bring the most comprehensive information into meta-analysis.

## 4. Discussion

In this study, we evaluated four different strategies handling the missingness of individual lifestyle components in the meta-analysis of gene-lifestyle interaction using LRS-stratified summary statistics from participating cohorts. We aimed to find the best way to leverage the available data while appropriately handling the heterogeneity due to missing data in the LRS, and further improve the power of identifying novel loci for the trait of interest. Only utilizing data contributed by the cohorts without missingness in any lifestyle components (the Complete Approach) is very underpowered, while freely meta-analyzing all the association results contributed by the cohorts even with missing components in the LRS (the Naïve Approach) is slightly liberal. The Safe Approach and Moderator Approach are both slightly conservative and their p-values are almost identical to each other. We also observed that, as expected, the more cohorts with information for all lifestyle components we used in meta-analysis, the more accurate the results. This result confirms our primary hypothesis.

A risk score is a commonly used approach to evaluate combined effects of risk factors and it may play an important role in personalized medicine. In the past, the scientific community has proposed several well-known risk scores. For example, the Framingham Risk Score (Wilson et al., 1998) is a sex-specific score used to estimate the 10-year cardiovascular risk, and the diabetes risk score (Lindström & Tuomilehto, 2003) is a screening tool for identifying subjects at high-risk for type 2 diabetes. Lifestyle Risk Scores have also become popular as people are increasingly interested in their clinical implications drawn by the joint effects of individual lifestyle factors to a specific trait, disease, or time-to event outcome. In the meantime, the genetic risk score (GRS) has become a widely used tool to improve identification of persons who are at risk for common complex diseases after numerous stories about exceptional success in genome-wide association studies (GWAS).

There have been some prior studies combining genetic risk scores and lifestyle risk scores to explore their joint behavior on risk of CVD (Abdullah Said et al., 2018) and Colorectal Cancer (Cho et al., 2019). Specifically, these studies divided study samples into subgroups based on the combination of genetic risk score level and lifestyle risk score level, and found that within and across genetic risk groups, adherence to poor behavioral lifestyle was associated with increased risk of diseases, and there was no interaction effect between genetic risk and lifestyle risk. This might seem discouraging whether adding genetic information could add much to the risk prediction studies using lifestyle risk scores. However, it is important to note that the genetic risk score was calculated based on variants reported from previous standard genome-wide significant analyses without taking its potential modification effect into consideration; variants whose effects may differ by level of lifestyle risk score might therefore be missed by standard GWAS screening. Moreover, a LRS may have a different modification effect on each variant, so instead of looking at aggregated genetic risk score only, interaction with one variant at a time should also be evaluated. Our study looked into the combination of genetic and lifestyle information in the way of performing meta-analysis of gene-by-lifestyle interaction in order to find novel loci for complex disease traits, and those potential novel loci may provide additional information for computing a genetic risk score, which could increase the power of previous studies.

Handling missing data in the aggregation of risk factors is challenging, yet important and worth the effort to explore in further detail. Based on the properties of genetic architecture, GRS can be computed using imputed or proxy SNPs, when the originally reported variants are not available, based on the largely available reference panel, such as 1000 Genome Project (Auton et al., 2015). Thus, it is more flexible than LRS in terms of dealing with missingness. There were several methods proposed to impute phenotypes using the correlation structure between phenotypes, family structure or information from other cohorts (Chen, Peloso, & Dupuis, 2018; Dahl et al., 2016; Hormozdiari et al., 2016), but these methods rarely dealt with the case that one phenotype is completely unavailable for all the individuals in one particular cohort contributing to a large meta-analysis, which is what we encountered in our study. When considering using summary statistics in meta-analysis, a previous study (Loef & Walach, 2012) tried to deal with the issue of missingness by restricting the study sample to cohorts with at least three out of five lifestyle behaviors available, reducing sample size and thus power to a great extent, with the issue of heterogeneity unresolved. Our study proposes making the best use of the available data gathered from cohorts to obtain accurate combined effects of risk factors, thereby providing a novel perspective for risk score based meta-analysis in future research.

Our study examined the Moderator Approach, which is a novel approach of accounting for missingness via meta-regression in the gene-by-environment interaction field. Instead of performing stratum-specific meta-analyses and then evaluating the interaction, this approach can achieve the final goal in one step via meta-regression, with meta-analysis results of both exposure groups as input. However, due to the meta-regression setting, the Moderator Approach requires that the number of cohorts have GWAS results available for a SNP (which equals to 4 in our study) is greater than the number of predictors divided by two (which equals to [one main effect + one interaction effect + four missingness effects]/2 = 3 in our study). Therefore, it restricted the analyses to the SNPs existing in the GWAS results of all four cohorts, thereby eliminating a large number of SNPs from the analyses and possibly missed positive signals. On the other hand, the design matrix of the meta-regression model in the Moderator Approach should be treated with caution because in some cases of missingness pattern, the design matrix would suffer from multicollinearity and we could not successfully obtain the least square estimates. Since the Safe Approach can provide almost identical results as the Moderator Approach but does not have a restriction on the missingness pattern and the number of cohorts and predictors, we would recommend using the Safe Approach to handle missingness during meta-analysis. Potential future directions of our study would be to further investigate the Moderator approach and to evaluate the performance of Safe Approach and Moderator Approach under larger scale of meta-analyses.

### Strengths and Limitations

Our study has several important strengths. To our knowledge, this is the very first study to explore how to deal with missingness in individual lifestyle components in order to improve the power for identifying novel genetic loci for complex disease traits. Our study performed thorough comparisons between four meta-analysis approaches via various cohort mixture scenarios, thus providing comprehensive information for investigators to refer to.

Although this study has several strengths as an innovative work for dealing with missingness in gene-by-lifestyle interaction, it has some limitations. When calculating the “Partially Missing” LRS, we assigned the missingness pattern to each of the cohorts. Also, when performing the meta-analyses using a mixture of results obtained from cohort-specific analysis conducted with “Complete” LRS or “Partially Missing” LRS, although we considered various cohort mixture scenarios, we still were not able to catch every possible pattern. This kind of design may lose some flexibility and consequently fail to capture all the information during the comparison. Moreover, our study mainly evaluated the performance of different approaches in terms of joint effects instead of focusing on the interaction effect. We did not manage to capture a clear pattern when comparing the interaction effect between different meta-analysis approaches, due to the small sample size of our study. It is worth pursuing the comparison of the interaction effect itself among different approaches with a larger sample size, by incorporating more cohorts in our next step.

In summary, we evaluated four approaches of incorporating the missingness of lifestyle components in the meta-analysis of gene-by-lifestyle interaction. Based on our results, we generally recommend using the Safe Approach since it is straightforward to implement and yields non-inflated results. Handling the missingness of individual lifestyle components properly can efficiently increase statistical power of gene-by-lifestyle interaction analysis for identifying novel loci of complex traits.

## Acknowledgements

This project was largely supported by a grant from the U.S. National Heart, Lung, and Blood Institute (NHLBI), the National Institutes of Health, R01HL118305. The full list of acknowledgments appears in the Supplementary Note.

