## Supplementary Materials for "Lifestyle Risk Score for aggregating multiple lifestyle factors: Handling missingness of individual lifestyle components in meta-analysis of gene-by-lifestyle interactions"

[*] Shared-first author

[#] Shared-last author

1) Department of Biostatistics, Boston University School of Public Health, Boston, MA, USA
2) Division of Statistical Genomics, Department of Genetics, Washington University School of Medicine, St. Louis, MO, USA
3) Human Genetics Center, Department of Epidemiology, Human Genetics, and Environmental Sciences, School of Public Health, the University of Texas School of Public health, Houston, TX, USA
4) Department of Human Genetics, Leiden University Medical Center, Leiden, the Netherlands
5) Field and Environmental Data Science, Benson Hill Inc, St. Louis, MO, USA
6) The Human Genome Sequencing Center, Baylor College of Medicine, Houston, Texas
7) NHLBI and Boston University Framingham Heart Study, Framingham, MA
8) Department of Medicine, Cardiovascular Division, Washington University School of Medicine, St. Louis, MO, USA
9) Section of Gerontology and Geriatrics, Department of Internal Medicine, Leiden University Medical Center, Leiden, the Netherlands
10) Department of Population Health Sciences, Duke University, Durham, NC, USA
11) Division of Endocrinology, Department of Internal Medicine, Leiden University Medical Center, Leiden, the Netherlands
12) Leiden Laboratory for Experimental Vascular Medicine, Leiden University Medical Center, Leiden, the Netherlands
13) Department of Psychiatry, Washington University School of Medicine, St. Louis, MO, USA
14) Division of Biostatistics, Washington University School of Medicine, St. Louis, MO, USA

### Supplementary Materials

**Supplementary Method……………………………………………………………..…..1**

**Supplementary Tables……………………………………………………………...5**

**Supplementary Figures…………………………………………………………...11**

**Supplementary Note..……………………………………………………..…...21**

### Supplementary Method: Moderator Approach

#### Adjusting for heterogeneity in GxE GWAS meta-analysis due to missingness in components of aggregated environmental variables

RJ Waken

April 25, 2019

### 1 Methods

#### 1.1 Motivating Example

To motivate our effort, we examine the case of a cardiovascular trait GxE GWAS using a lifestyle risk score (LRS) as the environmental term. We define a cutoff $k$ such that any individual with LRS $< k$ is in the unexposed group, and has a "heart-healthy" lifestyle, while any individual with LRS $\geq k$ is in the exposed group, and does not have a "heart-healthy" lifestyle. We can then combine summary statistics created in these individual strata using the stratified GxE meta-analysis model (commonly referred to as "Model 2")

$$\begin{aligned} \mathrm{E}\left[Y_{\mathrm{LRS}<k}\right] =& \boldsymbol{\beta}_{\mathrm{cov}} \times \mathrm{Covariates} + \beta_{\mathrm{Unexposed}} \times \mathrm{SNP} \\ \mathrm{E}\left[Y_{\mathrm{LRS}\geq k}\right] =& \boldsymbol{\beta}_{\mathrm{cov}} \times \mathrm{Covariates} + \beta_{\mathrm{Exposed}} \times \mathrm{SNP}, \end{aligned} \tag{1}$$

to assess the G and GxE effects in individual cohorts before meta-analysis.

##### 1.1.1 Missingness in aggregate environmental terms

Unfortunately, this GxLRS GWAS requires that all elements used in the LRS are present in a given cohort to appropriately stratify the individuals in any that cohort before summary statistics are fit through 1. When this is not the case, two options are currently considered:

- cohorts missing elements of the LRS will be omitted, which reduces the overall sample size and may induce selection bias
- cohorts missing elements of the LRS may inappropriately classify "exposed" individuals as "unexposed" before fitting the summary statistics required for 1, which leads to within stratum phenotypic heterogeneity in cases where GxLRS interactions are present

In either of the above cases, we expect this issue to result in a loss of power to detect significant G or GxE terms using 1.

#### 1.2 Proposal

We perform meta-analysis through the model

$$\hat{\boldsymbol{\beta}} = \boldsymbol{X}\boldsymbol{\alpha} + \boldsymbol{\varepsilon}, \tag{2}$$

where $\boldsymbol{X}$ is the meta-analysis design matrix, $\boldsymbol{\alpha}$ are the parameters of interest, typically $\boldsymbol{\varepsilon} \sim \text{MVN}(\mathbf{0}, \boldsymbol{V})$, $\hat{\boldsymbol{\beta}}$ are the summary G terms from 1, and $\boldsymbol{V}$ is the covariance matrix constructed from the summary statistics contributed by each of the cohorts. The $i^{th}$ row of our meta-analysis design matrix is $X_i = [I_{i\in\text{Unexposed}}, I_{i\in\text{Exposed}}]$, where $I$ is the indicator function. This parameterization is chosen for convenience, and equivalent to implementing meta-analysis through Easystrata. The hypothesis tests commonly used in this context are

$$H_0\colon \alpha_1 = \alpha_2 = 0,$$
$$H_1\colon \text{Not } H_0,$$

commonly referred to as the joint 2df test, and

$$H_0\colon \alpha_1 - \alpha_2 = 0,$$
$$H_1\colon \alpha_1 - \alpha_2 \neq 0,$$

commonly referred to as the interaction test.

Further, in our particular case, we can only misclassify individuals that belong in the "exposed" group into the "unexposed" in any given cohort that is missing elements of the aggregate environmental index. As a result, we only need to address misclassification in the situation where our summary statistics $i \in$ Unexposed. Although the current application only concerns heterogeneity in the unexposed group, the below method can be extended to the case where within exposure heterogeneity is possible in both groups.

#### 1.3 Moderator meta-analysis

We propose the use of moderator variables to account for the effect of missing elements of our aggregate environmental variable. This approach is attractive due to its simplicity and ease of implementation, but requires the assumption that the effect of missingness in the environmental term is independent of the resulting errors in meta-analysis, conditional on the indicator variable describing missingness. Our moderator meta-analysis model is

$$\hat{\boldsymbol{\beta}} = [\boldsymbol{X} : \boldsymbol{Z}]\,[\boldsymbol{\alpha} : \boldsymbol{\alpha_Z}] + \boldsymbol{\varepsilon}, \tag{3}$$

where $\boldsymbol{Z}$ are columns added to the design matrix that describe the incidence of missingness in the aggregated environmental term, and $\boldsymbol{\alpha_Z}$ are the parameters that describe the effect of missingness. Note that, in addition to the presented assumptions, this requires enough contributing cohorts to appropriately estimate $\boldsymbol{\alpha_Z}$. As a result of our special case, $Z_i$ will only have non-zero entries in cases where row $i$ represents summary statistics for the "unexposed" group.

##### 1.4 Obtaining estimates for $\boldsymbol{\alpha}$ and $\boldsymbol{\alpha}:\boldsymbol{\alpha_Z}$ in inverse variance weighted meta-analysis

Both formulations above in ((2)) and ((3)) can be modified to accommodate an inverse variance weighting scheme for meta analysis. Here, we find estimates for $\boldsymbol{\alpha}$ using

$$\hat{\boldsymbol{\alpha}} = \left(\boldsymbol{X}^T\boldsymbol{V}^{-1}\boldsymbol{X}\right)^{-1}\boldsymbol{X}^T\boldsymbol{V}^{-1}\hat{\boldsymbol{\beta}} \tag{4}$$

for (2), and

$$\hat{\boldsymbol{\alpha}}:\hat{\boldsymbol{\alpha}}_{\boldsymbol{Z}} = \left([\boldsymbol{X}:\boldsymbol{Z}]^T\boldsymbol{V}^{-1}[\boldsymbol{X}:\boldsymbol{Z}]\right)^{-1}[\boldsymbol{X}:\boldsymbol{Z}]^T\boldsymbol{V}^{-1}\hat{\boldsymbol{\beta}} \tag{5}$$

for (3), where $\boldsymbol{V}$ is the diagonal matrix of variances and $V_{ii} = \text{Var}\left(\hat{\beta}_i\right)$.

##### 1.5 Obtaining test statistics for the joint 2df and interaction tests

###### 1.5.1 Stratified meta-analysis with no moderator variables

The covariance matrix for (4) is $\left(\boldsymbol{X}^T\boldsymbol{V}^{-1}\boldsymbol{X}\right)^{-1}$. Note that, in the case of "Model 2," where main effects models have been fit in exposed and unexposed strata, $\left(\boldsymbol{X}^T\boldsymbol{V}^{-1}\boldsymbol{X}\right)^{-1}$ is diagonal.

In the case where $\boldsymbol{\alpha} = \boldsymbol{0}$,

$$\hat{\boldsymbol{\alpha}}^T\boldsymbol{X}^T\boldsymbol{V}^{-1}\boldsymbol{X}\hat{\boldsymbol{\alpha}} \sim \chi_2,$$

which yields the Wald type joint 2df test.

In the case where $[-1,\, 1]^T\,\boldsymbol{\alpha} = 0$,

$$\left(\begin{bmatrix}-1\\1\end{bmatrix}\hat{\boldsymbol{\alpha}}\right)^T\left(\begin{bmatrix}-1\\1\end{bmatrix}^T\boldsymbol{X}^T\boldsymbol{V}^{-1}\boldsymbol{X}^{-1}\begin{bmatrix}-1\\1\end{bmatrix}\right)^{-1}\hat{\boldsymbol{\alpha}}\begin{bmatrix}-1\\1\end{bmatrix} \sim \chi_1,$$

which yields the Wald type interaction test.

###### 1.5.2 Stratified meta-analysis with moderator variables

The covariance matrix for (5) is $\left([\boldsymbol{X}:\boldsymbol{Z}]^T\boldsymbol{V}^{-1}[\boldsymbol{X}:\boldsymbol{Z}]\right)^{-1}$. Note that we are not interested in the moderator variables. Let the covariance matrix for the estimate vector of interest $\hat{\boldsymbol{\alpha}}$ be $\hat{\Sigma}_{\hat{\boldsymbol{\alpha}}}$, which is the subset of $\left([\boldsymbol{X}:\boldsymbol{Z}]^T\boldsymbol{V}^{-1}[\boldsymbol{X}:\boldsymbol{Z}]\right)^{-1}$ whose rows and columns correspond to the $\hat{\boldsymbol{\alpha}}$ estimates in $\hat{\boldsymbol{\alpha}}:\hat{\boldsymbol{\alpha}}_{\boldsymbol{Z}}$. Note that $\hat{\Sigma}_{\hat{\boldsymbol{\alpha}}}$ is **NOT** diagonal.

In the case where $\boldsymbol{\alpha} = \boldsymbol{0}$,

$$\hat{\boldsymbol{\alpha}}^T\hat{\Sigma}_{\hat{\boldsymbol{\alpha}}}^{-1}\hat{\boldsymbol{\alpha}} \sim \chi_2,$$

which yields the Wald type joint 2df test.

In the case where $[-1,\, 1]^T \boldsymbol{\alpha} = 0$,

$$\left( \begin{bmatrix} -1 \\ 1 \end{bmatrix} \hat{\boldsymbol{\alpha}} \right)^T \left( \begin{bmatrix} -1 \\ 1 \end{bmatrix}^T \hat{\Sigma}_{\hat{\boldsymbol{\alpha}}} \begin{bmatrix} -1 \\ 1 \end{bmatrix} \right)^{-1} \hat{\boldsymbol{\alpha}} \begin{bmatrix} -1 \\ 1 \end{bmatrix} \sim \chi_1,$$

which yields the Wald type interaction test.

**Supplementary Table S1:** Genotyping, Imputations and analyses software

| Study Name | Ancestry | Study Design [1] | Which PCs? | Other Covariates [2] | Genotyping Platforms | Genotyping Calling Algorithm |
|---|---|---|---|---|---|---|
| **ARIC** | EA | UN | 1-10 | Field centers | Affymetrix Genome-Wide Human SNP Array 6.0 | Birdseed calling algorithm |
| **FHS** | EA | FB | PC1-PC10 | cohort | Affymetrix Nsp, Sty and 50K gene centric | Birdseed |
| **HyperGen** | EA | FB | 0 | N/A | Affymetrix 5.0 | BRLMM |
| **NEO** | EA | UN | 4 | N/A | Illumina HumanCoreExome-24v1_A Beadchip | Gencall |

| Study Name | Imputation Software | Imputation Reference Panel | Build | Analysis Software | Robust or model-based statistics? | Family Studies: Method of handling relatedness WITHIN STRATA |
|---|---|---|---|---|---|---|
| **ARIC** | IMPUTE2 | 1000G Phase I [2010-11] - European panel (CEU, TSI, GBR, FIN, IBS) | hg19/b37 | MMAP | Robust | N/A |
| **FHS** | MaCH (v. 1.0.16) | 1000G Phase I v3 Shapeit2 Reference (2010-11 data freeze, 2013-09 haplotypes). ALL reference panel-- no monomorphic sites. | hg19 | MMAP | Robust | Kinship |
| **HyperGen** | MaCH/minimac | 1000G Phase 1 Integrated Release Version 3 Haplotypes (2010-11 data freeze, 2012-03-14 haplotypes) on 1,092 individuals of all ethnic backgrounds and excludes monomorphic and singleton sites | hg19/b36 | MMAP | Robust | Kinship |
| **NEO** | IMPUTE2 | 2011 v3 | hg19/b36 | ProBABEL | Robust | N/A |

[1] Cohort unrelated (UN), family-based (FB), case-control (CC)

[2] Other than those specified in the replication analysis plan.

**Supplementary Table S2:** General study characteristics

| Study Name | Ancestry | N | % Male | % Using Anti-hypertension Medicine |
|---|---|---|---|---|
| **ARIC** | EA | 9426 | 46.9 | 25.4 |
| **FHS** | EA | 7638 | 45.7 | 7.8 |
| **HyperGen** | EA | 1249 | 50.0 | 50.7 |
| **NEO** | EA | 5735 | 48.1 | 44.3 |

| Study Name | Age | | | | |
|---|---|---|---|---|---|
| | **Mean** | **SD** | **Median** | **Min.** | **Max.** |
| **ARIC** | 54.3 | 5.7 | 54.0 | 44.0 | 66.0 |
| **FHS** | 42.0 | 8.9 | 42.0 | 19.0 | 72.0 |
| **HyperGen** | 49.7 | 13.9 | 49.0 | 18.0 | 80.0 |
| **NEO** | 57.0 | 6.0 | 56.0 | 44.0 | 66.0 |

| Study Name | BMI | | | | |
|---|---|---|---|---|---|
| | **Mean** | **SD** | **Median** | **Min.** | **Max.** |
| **ARIC** | 27.0 | 4.9 | 26.3 | 14.4 | 56.3 |
| **FHS** | 26.2 | 5.0 | 25.4 | 15.4 | 60.5 |
| **HyperGen** | 29.3 | 5.9 | 28.5 | 15.1 | 64.5 |
| **NEO** | 30.0 | 4.8 | 29.5 | 17.2 | 57.9 |

**Supplementary Table S3:** Descriptive statistics of Systolic blood pressure, splitting by exposure group

| Study Name | All participants included in analysis | | | | | |
|---|---|---|---|---|---|---|
| | N | Mean | SD | Median | Min. | Max. |
| **ARIC** | 9426 | 122.2 | 19.4 | 120.0 | 61.0 | 221.0 |
| **FHS** | 7637 | 120.0 | 16.4 | 118.0 | 70.0 | 208.0 |
| **HyperGen** | 1249 | 130.9 | 23.1 | 129.0 | 79.5 | 231.0 |
| **NEO** | 5735 | 137.5 | 19.4 | 136.0 | 84.7 | 228.3 |

| Study Name | DLRS-C* = 0 | | | | | |
|---|---|---|---|---|---|---|
| | N | Mean | SD | Median | Min. | Max. |
| **ARIC** | 1966 | 119.5 | 17.8 | 117.0 | 72.0 | 197.0 |
| **FHS** | 2178 | 116.7 | 14.7 | 115.0 | 84.0 | 195.0 |
| **HyperGen** | 292 | 122.5 | 19.7 | 119.5 | 79.5 | 176.5 |
| **NEO** | 179 | 132.6 | 19.2 | 131.0 | 96.0 | 195.0 |

| Study Name | DLRS-C = 1 | | | | | |
|---|---|---|---|---|---|---|
| | N | Mean | SD | Median | Min. | Max. |
| **ARIC** | 7460 | 122.9 | 19.8 | 120.0 | 61.0 | 221.0 |
| **FHS** | 5459 | 121.4 | 16.8 | 119.0 | 70.0 | 208.0 |
| **HyperGen** | 957 | 133.5 | 23.5 | 131.5 | 82.0 | 231.0 |
| **NEO** | 5556 | 137.6 | 19.3 | 136.0 | 84.7 | 228.3 |

| Study Name | DLRS-M** = 0 | | | | | |
|---|---|---|---|---|---|---|
| | N | Mean | SD | Median | Min. | Max. |
| ARIC | 3983 | 121.5 | 18.8 | 119.0 | 72.0 | 207.0 |
| FHS | 3252 | 116.9 | 14.9 | 115.0 | 70.0 | 195.0 |
| HyperGen | 426 | 125.1 | 21.4 | 122.0 | 79.5 | 208.0 |
| NEO | 1121 | 135.0 | 18.9 | 133.3 | 90.0 | 210.7 |

| Study Name | DLRS-M = 1 | | | | | |
|---|---|---|---|---|---|---|
| | N | Mean | SD | Median | Min. | Max. |
| ARIC | 5443 | 122.7 | 19.9 | 120.0 | 61.0 | 221.0 |
| FHS | 4385 | 122.3 | 17.0 | 120.0 | 82.0 | 208.0 |
| HyperGen | 823 | 133.9 | 23.5 | 131.5 | 82.0 | 231.0 |
| NEO | 4614 | 138.1 | 19.4 | 136.3 | 84.7 | 228.3 |

*DLRS-C = Dichotomous LRS with Complete Components

**DLRS-M = Dichotomous LRS with Partially Missing Components

**Supplementary Table S4:** Sample distribution on the combination of DLRS-C* and DLRS-M**

| | (N's) | | DLRS-M | | |
|---|---|---|---|---|---|
| | | | **E0*** ** | **E1** | ***Total*** |
| **ARIC** | **DLRS-C** | **E0** | 1966 | 0 | 1966 |
| | | **E1** | 2017 | 5443 | 7460 |
| | | ***Total*** | 3983 | 5443 | 9426 |
| **FHS** | **(N's)** | | **DLRS-M** | | |
| | | | **E0** | **E1** | ***Total*** |
| | **DLRS-C** | **E0** | 2178 | 0 | 2178 |
| | | **E1** | 1074 | 4386 | 5460 |
| | | ***Total*** | 3252 | 4386 | 7638 |
| **HyperGen** | **(N's)** | | **DLRS-M** | | |
| | | | **E0** | **E1** | ***Total*** |
| | **DLRS-C** | **E0** | 292 | 0 | 292 |
| | | **E1** | 134 | 823 | 957 |
| | | ***Total*** | 426 | 823 | 1249 |
| **NEO** | **(N's)** | | **DLRS-M** | | |
| | | | **E0** | **E1** | ***Total*** |
| | **DLRS-C** | **E0** | 179 | 0 | 179 |
| | | **E1** | 942 | 4614 | 5556 |
| | | ***Total*** | 1121 | 4614 | 5735 |

*DLRS-C = Dichotomous LRS with Complete Components

**DLRS-M = Dichotomous LRS with Partially Missing Components

***E0 = unexposed group; E1 = exposed group

**Supplementary Figure S1:** Q-Q plot of different approaches compared to "benchmark" results for Scenario 2 (Scenario 2: Use "complete" results from FHS, and "partially Missing" results from HyperGEN, ARIC and NEO). $\lambda_{Naive}$ = 1.007, $\lambda_{Safe}$ = $\lambda_{Moderator}$ =0.993, $\lambda_{Complete}$ = 0.986.

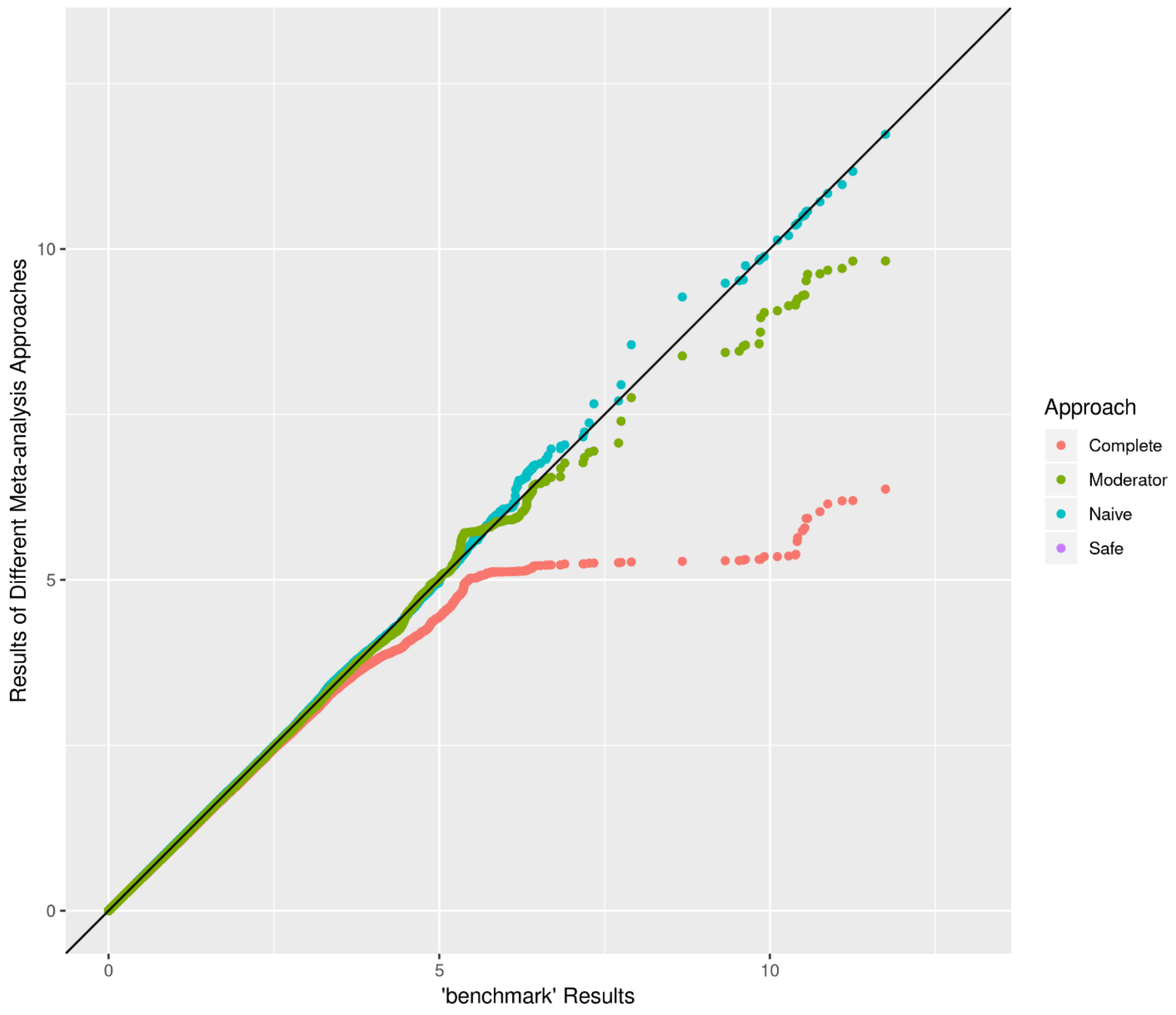

**Supplementary Figure S2:** Q-Q plot of different approaches compared to "benchmark" results for Scenario 3 (Scenario 3: Use "complete" results from ARIC and NEO, and "partially Missing" results from HyperGEN, FHS). $\lambda_{Naive} = 1.002$, $\lambda_{Safe} = \lambda_{Moderator} = 0.989$, $\lambda_{Complete} = 0.978$.

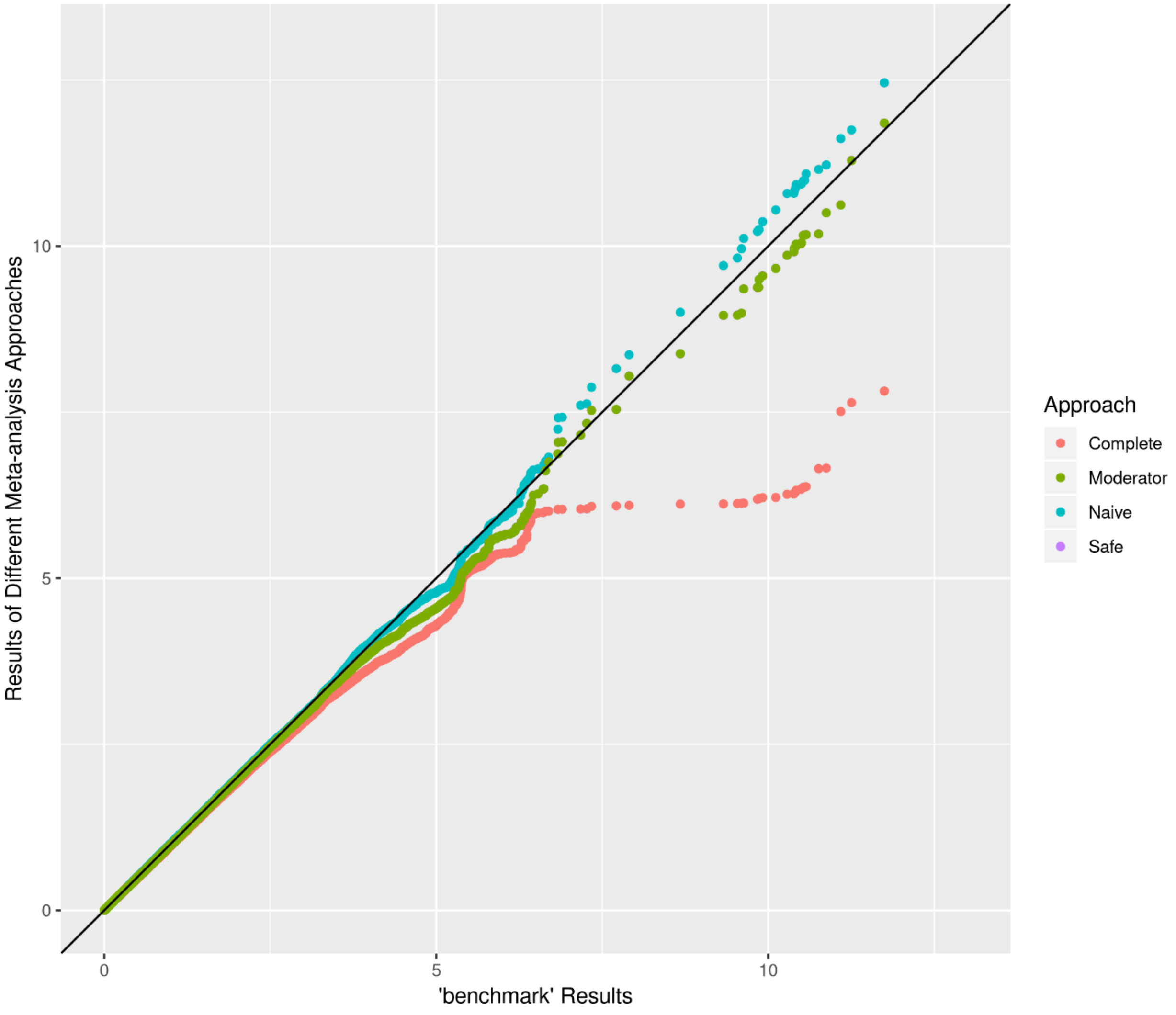

**Supplementary Figure S3:** Q-Q plot of different approaches compared to "benchmark" results for Scenario 4 (Scenario 4: Use "complete" results from ARIC, HyperGEN and NEO, and "partially Missing" results from FHS). $\lambda_{Naive} = 1.005$, $\lambda_{Safe} = \lambda_{Moderator} = 0.997$, $\lambda_{Complete} = 0.991$.

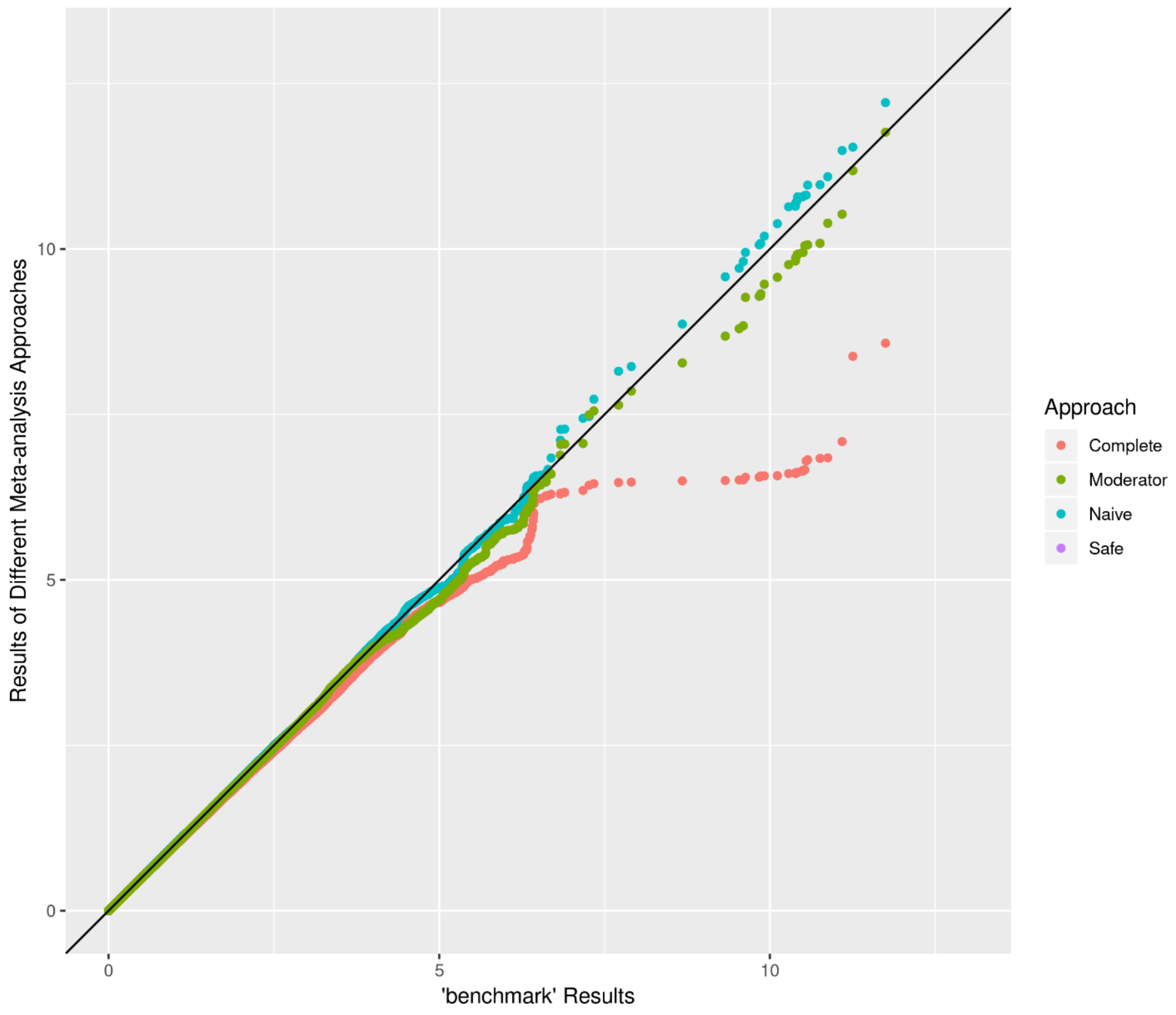

**Supplementary Figure S4:** Scatterplots of comparison of four approaches to "benchmark" results in terms of $-\log_{10}$ (p-value) for Scenario 2. Each plot shows SNPs with p-value$<10^{-6}$ for any of the two approaches being compared in the plot. SNPs reaching genome-wide significant (p-value$<5\times10^{-8}$) in "benchmark" results are marked as triangle. (Scenario 2: Use "complete" results from FHS, and "partially Missing" results from HyperGEN, ARIC and NEO)

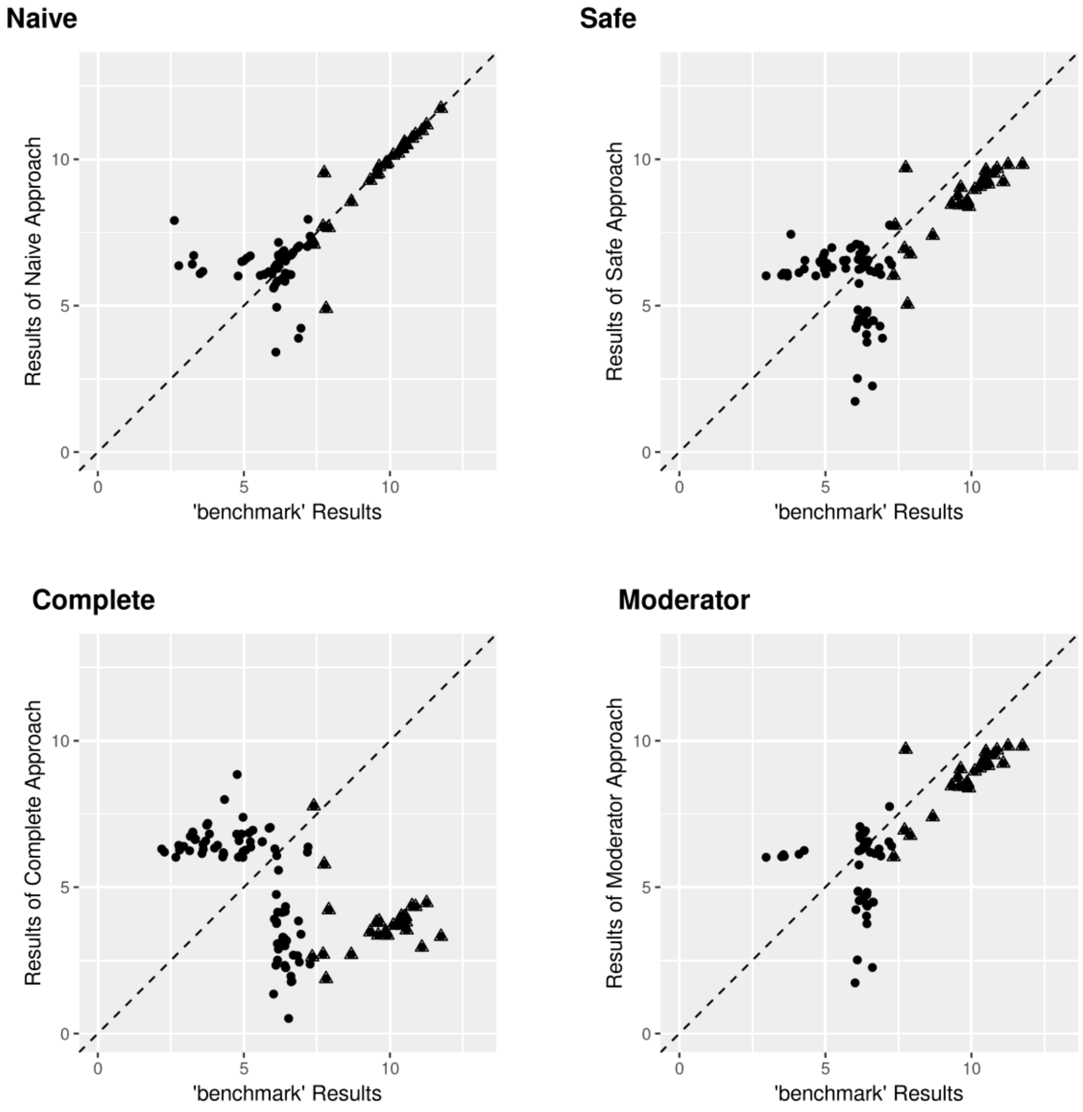

**Supplementary Figure S5:** Scatterplots of comparison between four approaches in terms of –$\log_{10}$ (p-value) for Scenario 2. Each plot shows SNPs with p-value<$10^{-6}$ for any of the two approaches being compared in the plot. (Scenario 2: Use "complete" results from FHS, and "partially Missing" results from HyperGEN, ARIC and NEO)

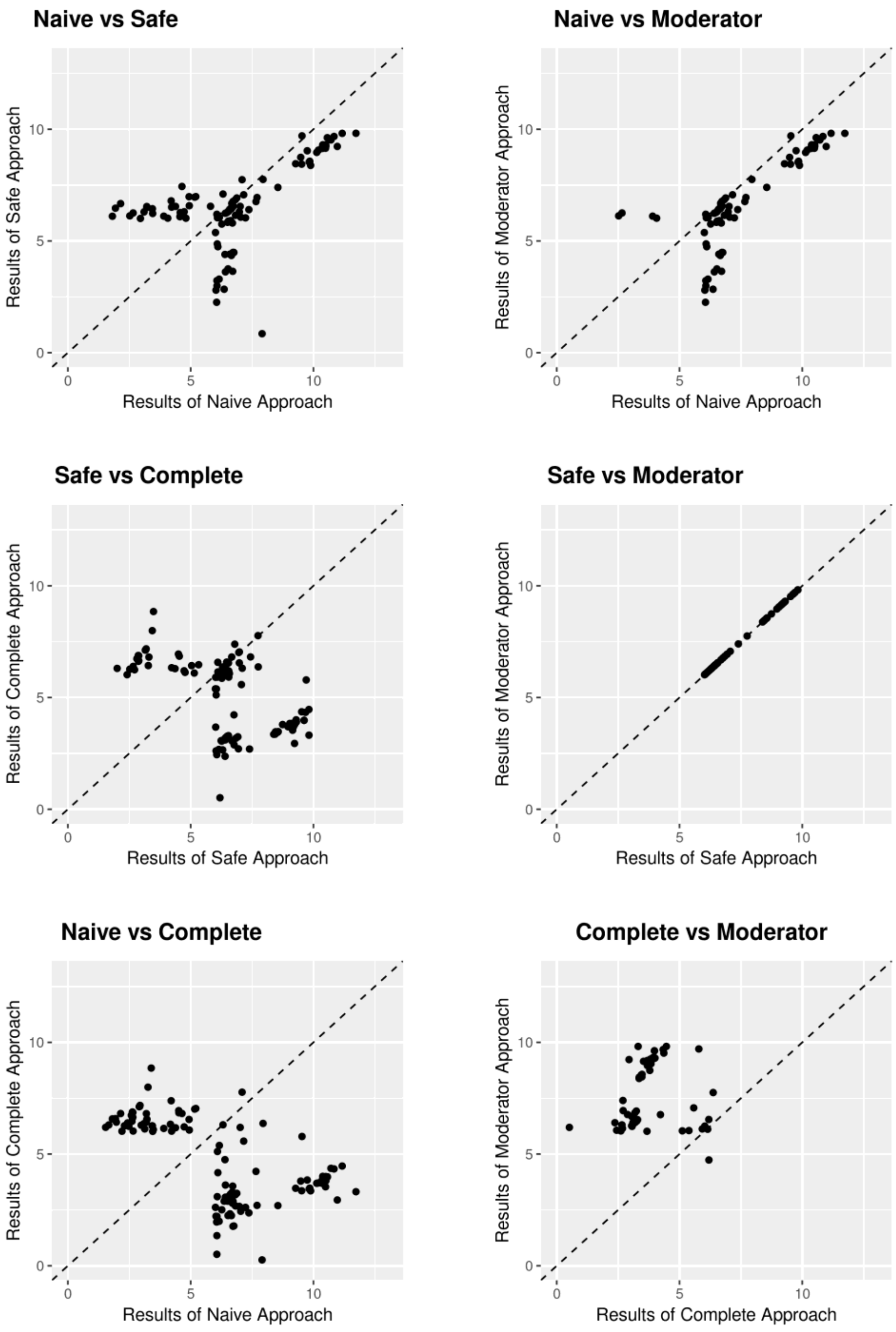

**Supplementary Figure S6:** Scatterplots of comparison of four approaches to "benchmark" results in terms of $-\log_{10}$ (p-value) for Scenario 3. Each plot shows SNPs with p-value<$10^{-6}$ for any of the two approaches being compared in the plot. SNPs reaching genome-wide significant (p-value<$5\times10^{-8}$) in "benchmark" results are marked as triangle. (Scenario 3: Use "complete" results from ARIC and NEO, and "partially Missing" results from HyperGEN, FHS)

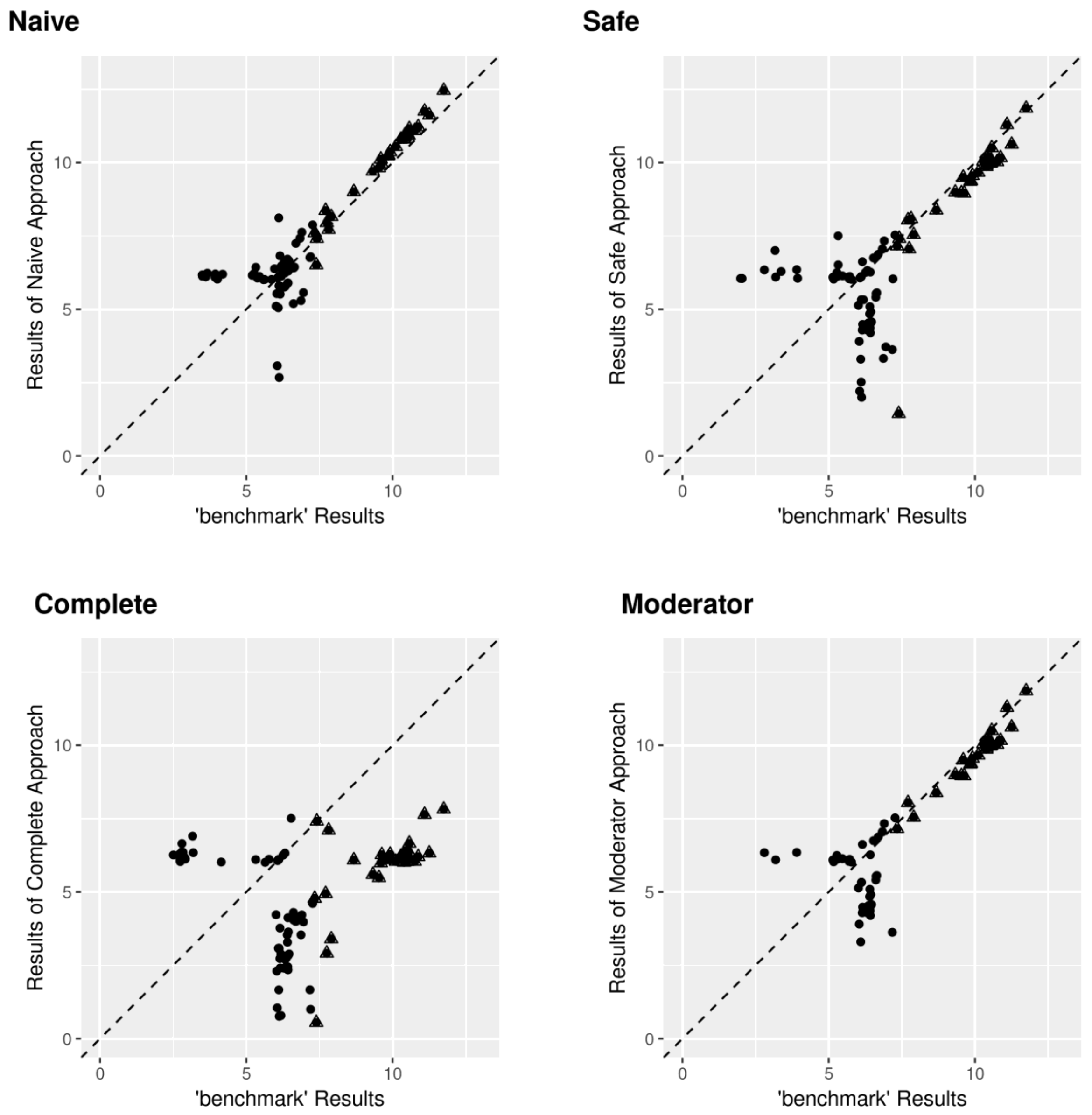

**Supplementary Figure S7:** Scatterplots of comparison between four approaches in terms of –$\log_{10}$ (p-value) for Scenario 3. Each plot shows SNPs with p-value<$10^{-6}$ for any of the two approaches being compared in the plot. (Scenario 3: Use "complete" results from ARIC and NEO, and "partially Missing" results from HyperGEN, FHS)

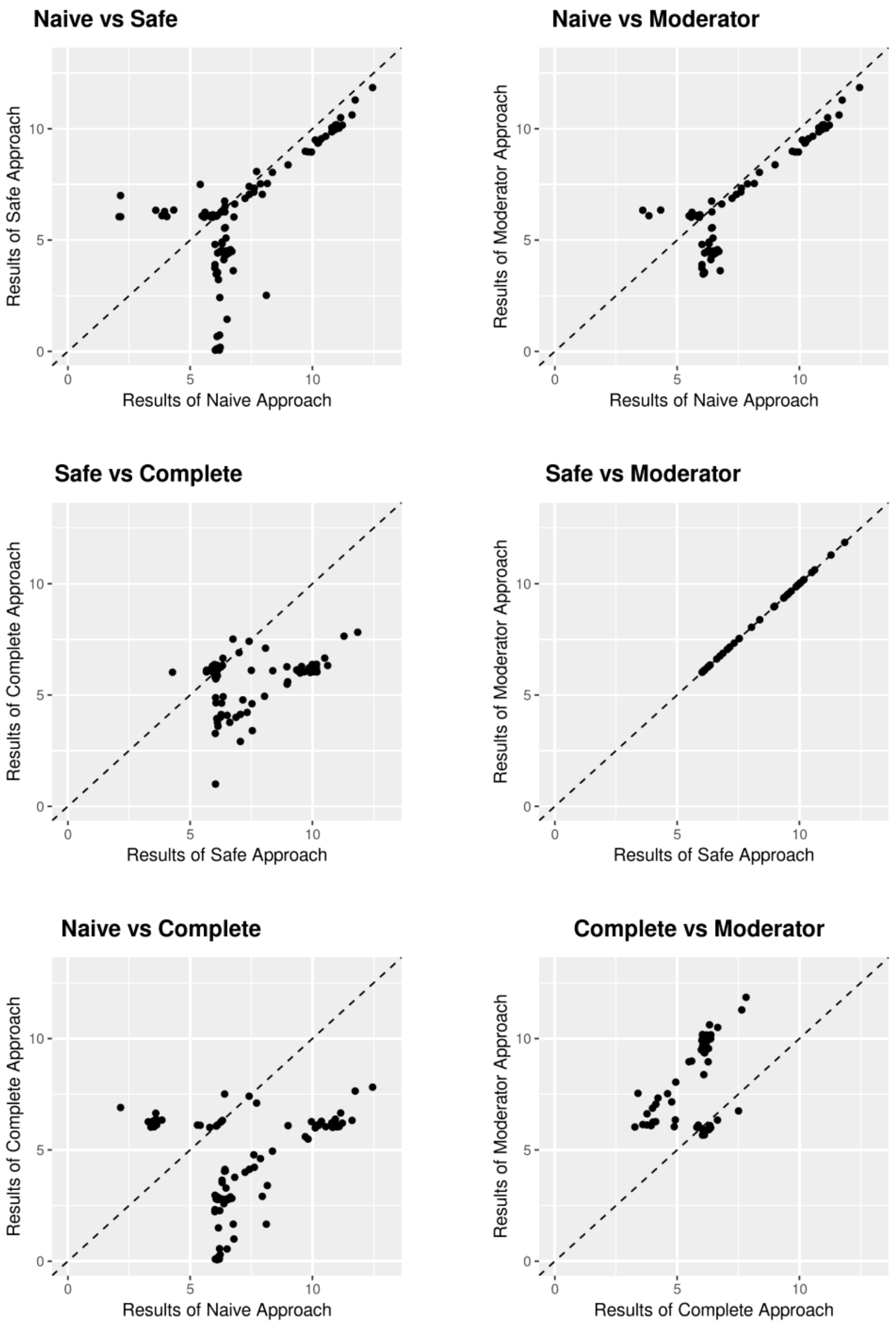

**Supplementary Figure S8:** Scatterplots of comparison of four approaches to "benchmark" results in terms of $-\log_{10}$ (p-value) for Scenario 4. Each plot shows SNPs with p-value<$10^{-6}$ for any of the two approaches being compared in the plot. SNPs reaching genome-wide significant (p-value<$5 \times 10^{-8}$) in "benchmark" results are marked as triangle. (Scenario 4: Use "complete" results from ARIC, HyperGEN and NEO, and "partially Missing" results from FHS)

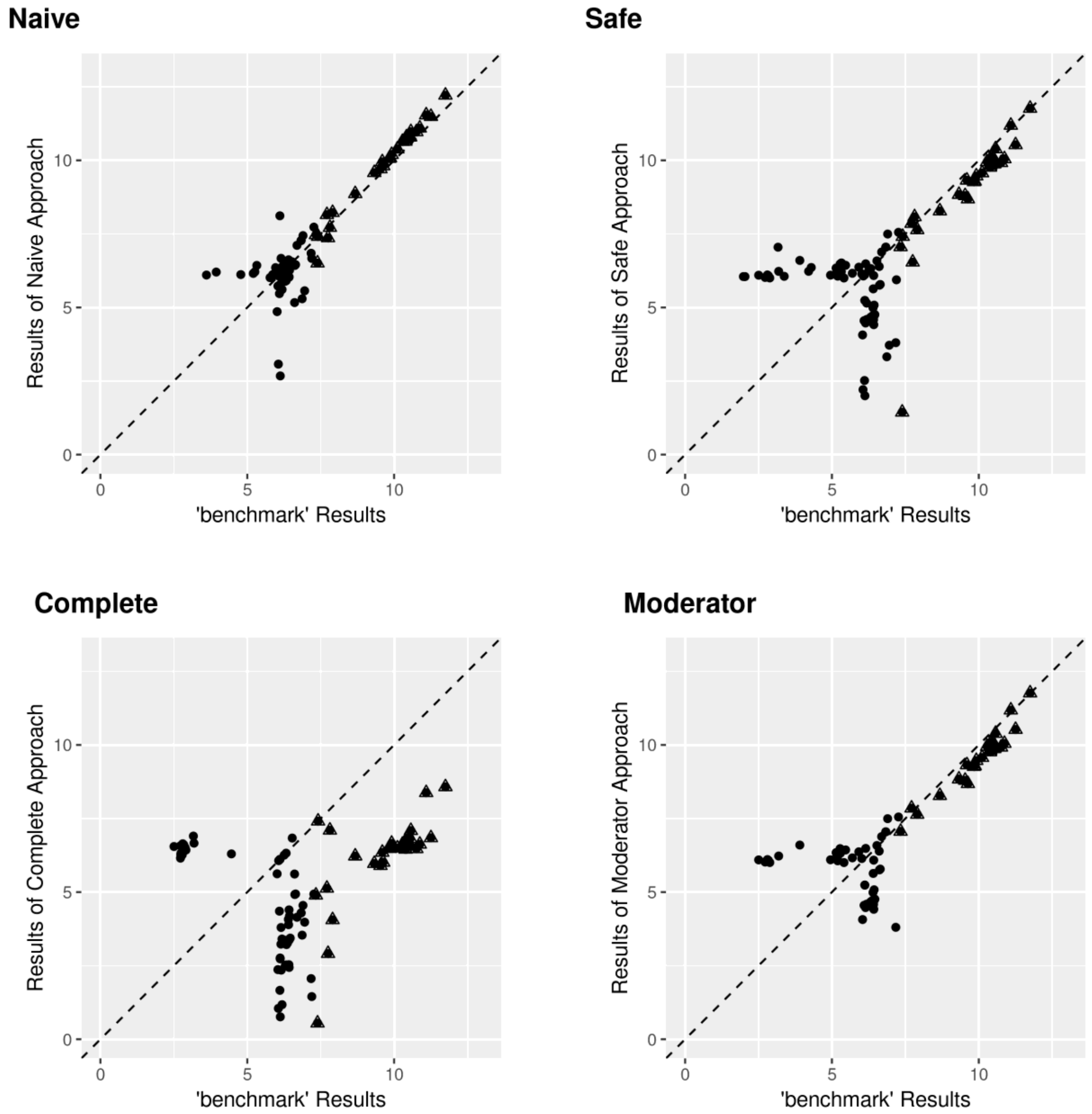

**Supplementary Figure S9:** Scatterplots of comparison between four approaches in terms of –$\log_{10}$ (p-value) for Scenario 4. Each plot shows SNPs with p-value<$10^{-6}$ for any of the two approaches being compared in the plot. (Scenario 4: Use "complete" results from ARIC, HyperGEN and NEO, and "partially Missing" results from FHS)

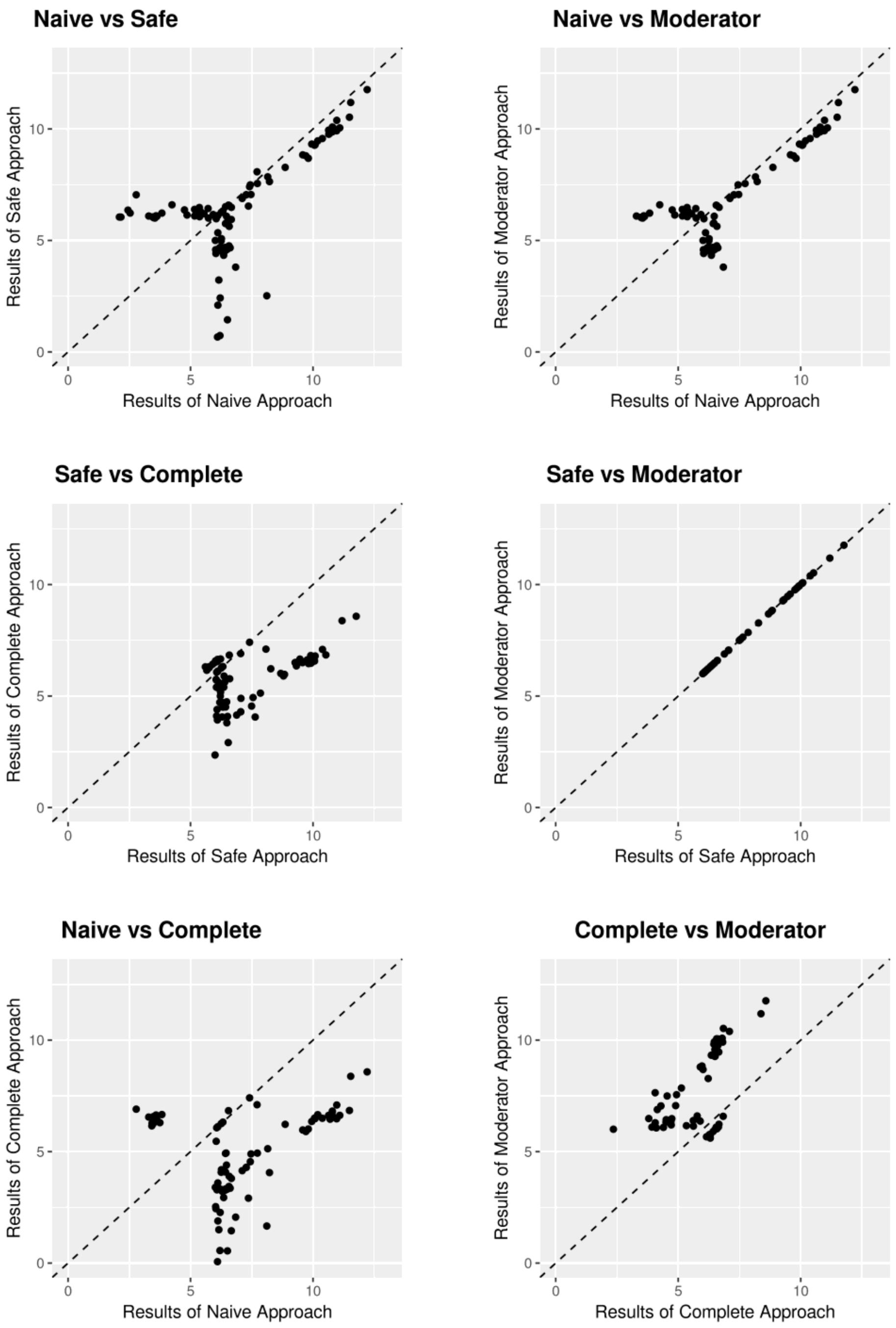

**Supplementary Figure S10:** Q-Q plot of different scenarios compared to “benchmark” results for Safe Approach. (Scenario 1: use “complete” results from ARIC, and “partially missing” results from HyperGEN, FHS and NEO; Scenario 2: Use “complete” results from FHS, and “partially Missing” results from HyperGEN, ARIC and NEO; Scenario 3: Use “complete” results from ARIC and NEO, and “partially Missing” results from HyperGEN, FHS; Scenario 4: Use “complete” results from ARIC, HyperGEN and NEO, and “partially Missing” results from FHS). $\lambda_{scenario\ 1}$=0.983, $\lambda_{scenario\ 2}$=0.994, $\lambda_{scenario\ 3}$=0.984, $\lambda_{scenario\ 4}$=0.991.

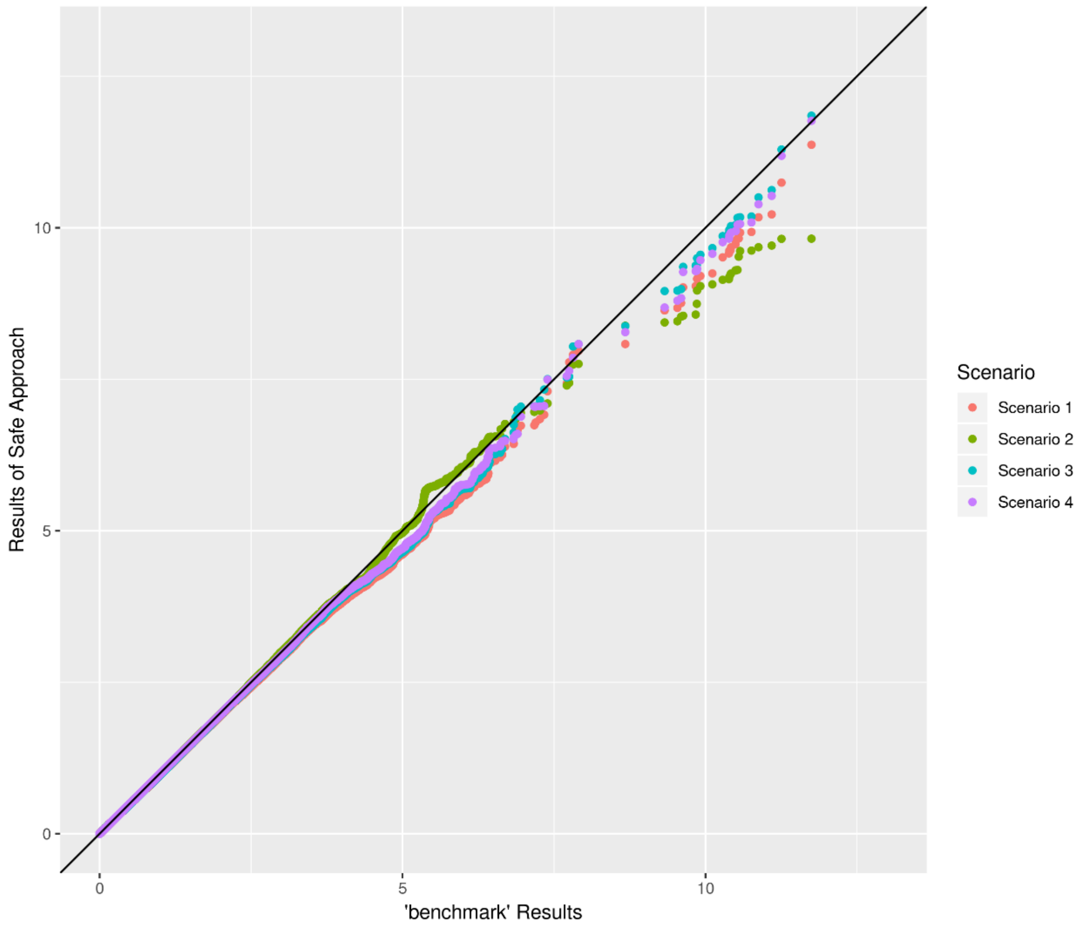

#### Supplementary Note: Study Acknowledgements

**ARIC (Atherosclerosis Risk in Communities) Study:** The ARIC study has been supported by the National Heart, Lung, and Blood Institute, National Institutes of Health, Department of Health and Human Services (contract numbers HHSN268201700001I, HHSN268201700002I, HHSN268201700003I, HHSN268201700004I and HHSN268201700005I), R01HL087641, R01HL059367 and R01HL086694; National Human Genome Research Institute contract U01HG004402; and National Institutes of Health contract HHSN268200625226C. Paul S. de Vries was supported by American Heart Association grant number 18CDA34110116.The authors thank the staff and participants of the ARIC study for their important contributions. Infrastructure was partly supported by Grant Number UL1RR025005, a component of the National Institutes of Health and NIH Roadmap for Medical Research.

**FHS (Framingham Heart Study):** This research was conducted in part using data and resources from the Framingham Heart Study of the National Heart Lung and Blood Institute of the National Institutes of Health and Boston University School of Medicine. The analyses reflect intellectual input and resource development from the Framingham Heart Study investigators participating in the SNP Health Association Resource (SHARe) project. This work was partially supported by the National Heart, Lung and Blood Institute's Framingham Heart Study (Contract Nos. N01-HC-25195 and HHSN268201500001I) and its contract with Affymetrix, Inc for genotyping services (Contract No. N02-HL-6-4278). A portion of this research utilized the Linux Cluster for Genetic Analysis (LinGA-II) funded by the Robert Dawson Evans Endowment of the Department of Medicine at Boston University School of Medicine and Boston Medical Center. This research was partially supported by grant R01- DK089256 from the National Institute of Diabetes and Digestive and Kidney Diseases (MPIs: Ingrid B. Borecki, L. Adrienne Cupples, Kari North).

**HyperGEN (Hypertension Genetic Epidemiology Network):** The Hypertension Network is funded by cooperative agreements (U10) with NHLBI: HL54471, HL54472, HL54473, HL54495, HL54496, HL54497, HL54509, HL54515, and 2 R01 HL55673-12. The study involves: University of Utah: (Network Coordinating Center, Field Center, and Molecular Genetics Lab); Univ. of Alabama at Birmingham: (Field Center and Echo Coordinating and Analysis Center); Medical College of Wisconsin: (Echo Genotyping Lab); Boston University: (Field Center); University of Minnesota: (Field Center and Biochemistry Lab); University of North Carolina: (Field Center); Washington University: (Data Coordinating Center); Weil Cornell Medical College: (Echo Reading Center); National Heart, Lung, & Blood Institute. For a complete list of HyperGEN Investigators: http://www.biostat.wustl.edu/hypergen/Acknowledge.html

**NEO (The Netherlands Epidemiology of Obesity study):** The authors of the NEO study thank all individuals who participated in the Netherlands Epidemiology in Obesity study, all participating general practitioners for inviting eligible participants and all research nurses for collection of the data. We thank the NEO study group, Petra Noordijk, Pat van Beelen and Ingeborg de Jonge for the coordination, lab and data management of the NEO study. The genotyping in the NEO study was supported by the Centre National de Génotypage (Paris, France), headed by Jean-Francois Deleuze. The NEO study is supported by the participating Departments, the Division and the Board of Directors of the Leiden University Medical Center, and by the Leiden University, Research Profile Area Vascular and Regenerative Medicine. Dennis Mook-Kanamori is supported by Dutch Science Organization (ZonMW-VENI Grant 916.14.023).
